# A mechanism that integrates microtubule motors of opposite polarity at the kinetochore corona

**DOI:** 10.1101/2023.04.25.538277

**Authors:** Verena Cmentowski, Giuseppe Ciossani, Ennio d’Amico, Sabine Wohlgemuth, Mikito Owa, Brian Dynlacht, Andrea Musacchio

## Abstract

Chromosome biorientation on the mitotic spindle is prerequisite to errorless genome inheritance. CENP-E (kinesin 7) and Dynein-Dynactin (DD), microtubule motors with opposite polarity, promote biorientation from the kinetochore corona, a polymeric structure whose assembly requires MPS1 kinase. The corona’s building block consists of ROD, Zwilch, ZW10, and the DD adaptor Spindly (RZZS). How CENP-E and DD are scaffolded and mutually coordinated in the corona remains unclear. Here, we report near-complete depletion of RZZS and DD from kinetochores after depletion of CENP-E and the outer kinetochore protein KNL1. With inhibited MPS1, CENP-E, which we show binds directly to RZZS, is required to retain kinetochore RZZS. An RZZS phosphomimetic mutant bypasses this requirement. With active MPS1, CENP-E is dispensable for corona expansion, but strictly required for physiological kinetochore accumulation of DD. Thus, we identify the corona as an integrated scaffold where CENP-E kinesin controls DD kinetochore loading for coordinated bidirectional transport of chromosome cargo.

---

As points of attachment of chromosomes to spindle microtubules in mitosis and meiosis, kinetochores are pivotal for chromosome segregation and genome inheritance (Cheeseman & Desai, 2008; Musacchio & Desai, 2017). Kinetochores are layered structures built on specialized chromosome loci named centromeres (Figure 1A). A centromere-proximal complex assembled on centromere landmarks and consisting of 16 subunits in humans (the constitutive centromere associated network or CCAN) recruits a centromere-distal layer involved in microtubule binding, the KMN network (Cheeseman & Desai, 2008). The KMN network forms from three constituent subcomplexes, the Knl1 complex (Knl1C, two subunits), the Mis12 complex (Mis12C, four subunits), and the Ndc80 complex (Ndc80C, four subunits). The Mis12C is a scaffold required for coordinated recruitment of the Knl1C and the Ndc80C (Cheeseman & Desai, 2008; Musacchio & Desai, 2017). Ndc80C is a microtubule receptor that promotes mature, ‘end-on’ interaction observed during metaphase, when spindle microtubules orient perpendicularly to the outer kinetochore (Cheeseman *et al*, 2006; DeLuca *et al*, 2006). Ndc80C and Knl1C also represent distinct but interconnected branches of an outer kinetochore regulatory network that controls bi-orientation and the timing of mitotic exit, as explained below.

**Figure 1.**
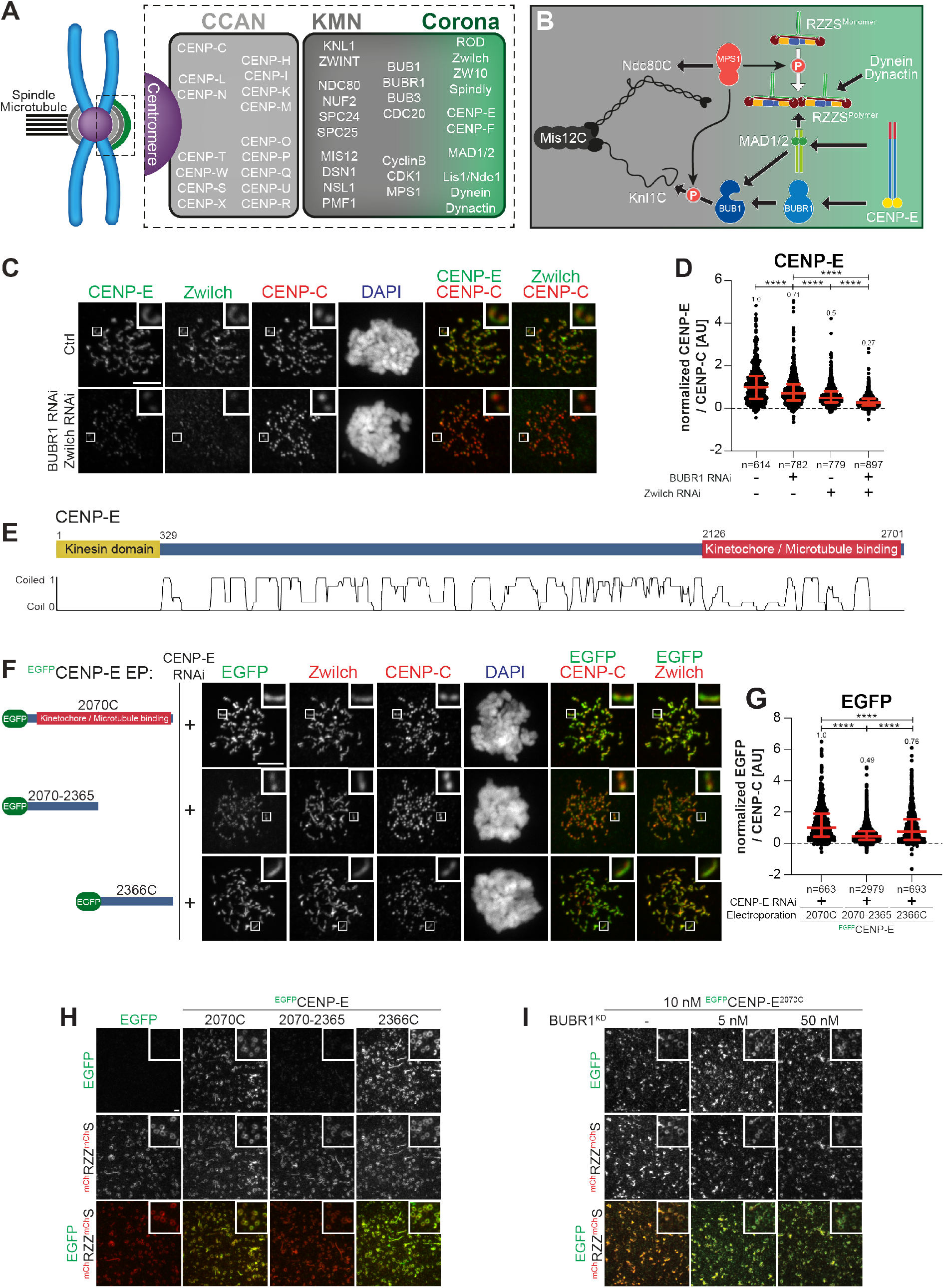
CENP-E^2070C^ contains binding sites for BUBR1 and RZZS. (**A**) Organization of the human kinetochore and corona. (**B**) Drawing depicting the hierarchical organization of outer kinetochore and kinetochore corona components. Thick arrows indicate recruitment of a protein to the protein indicated by the arrowhead. Thin arrows indicate phosphorylation. The white arrow indicates polymerization. (**C**) Representative images of the localization of CENP-E after depletion of Zwilch and BUBR1. Zwilch RNAi treatment was performed with 100 nM siRNA for 72 h. 48 h after Zwilch RNAi treatment cells were transfected with 100 nM BUBR1 siRNA. 8 h after transfection, cells were synchronized in G2 phase with 9 μM RO3306 for 15 h and then released into mitosis. Subsequently, cells were immediately treated with 3.3 μM Nocodazole, 10 μM MG132 for an additional hour. CENP-C was used to visualize kinetochores and DAPI to stain DNA. Scale bar: 5 μm. (**D**) Quantification of CENP-E levels at kinetochores of the experiment shown in panel C. n refers to individually measured kinetochores. (**E**) Organization of CENP-E with coiled-coil prediction. (**F**) Representative images showing the localization of different ^EGFP^CENP-E constructs in prometaphase after depletion of CENP-E with 60 nM siRNA. 13 h after RNAi treatment cells were electroporated with electroporation buffer or recombinant ^EGFP^CENP-E constructs as indicated. Following an 8 h recovery, cells were synchronized in G2 phase with 9 μM RO3306 for 15 h and then released into mitosis. Subsequently, cells were immediately treated with 3.3 μM Nocodazole for an additional hour. CENP-C was used to visualize kinetochores and DAPI to stain DNA. Scale bar: 5 μm. (**G**) Quantification of EGFP levels at kinetochores of the experiment shown in panel F. n refers to individually measured kinetochores. (**H**) ^mCH^RZZ^mCh^S ring-binding assays showing the recruitment of various ^EGFP^CENP-E truncations (10 nM concentration) to ^mCH^RZZ^mCh^S rings (approx. 40 nM concentration). Scale bar: 5 µm. (**I**) RZZS ring-binding assays showing the recruitment of ^EGFP^CENP-E^2070C^ to ^mCH^RZZ^mCh^S rings is unaffected by increasing concentrations of BUBR1^KD^. Scale bar: 5 µm.

In mitotic prometaphase, before the achievement of end-on binding and bi-orientation, outer kinetochores of metazoans assemble an outermost dense layer covered by fibrous material and named the kinetochore corona (Kops & Gassmann, 2020) (Figure 1A-B). The building block of the corona is a complex of the ROD-Zwilch-ZW10 (RZZ) complex with Spindly, a Dynein-Dynactin (DD) adaptor (Mosalaganti *et al*, 2017; Pereira *et al*, 2018; Raisch *et al*, 2022; Sacristan *et al*, 2018). MPS1, a protein kinase with various regulatory functions at mitotic kinetochores, phosphorylates ROD on two N-terminal residues to promote the polymerization of RZZ-Spindly (RZZS) that assembles the corona (Raisch *et al.*, 2022; Rodriguez-Rodriguez *et al*, 2018).

Two microtubule motors with opposite polarity facilitate chromosome alignment in prometaphase from the kinetochore corona by transporting chromosomes as cargoes while walking along the microtubule lattice. One is the homo-dimeric plus-end-directed kinesin-7 centromere protein E (CENP-E, 2701 residues in humans) (Yen *et al*, 1991; Yen *et al*, 1992). Like other kinesins, CENP-E is autoinhibited, likely through an interaction of its N- and C-terminal regions that is regulated by Aurora kinase phosphorylation (Craske *et al*, 2022; Espeut *et al*, 2008; Kim *et al*, 2008; Kim *et al*, 2010; Vitre *et al*, 2014). How CENP-E autoinhibition regulates end-on attachment, however, remains unclear.

The other motor in the kinetochore corona is the minus-end-directed Dynein (Gassmann, 2023; Pfarr *et al*, 1990), a 1.4 MDa complex composed of two copies each of six distinct polypeptides (Carter *et al*, 2016; Reck-Peterson *et al*, 2018). On its own, Dynein is poorly motile. Its motility is greatly enhanced by Dynactin, a 1.1 MDa 23-subunit assembly built from 11 distinct polypeptides. The interaction of Dynein and Dynactin is reinforced by specialized activating adaptor molecules (Carter *et al.*, 2016; Reck-Peterson *et al.*, 2018). Activating adaptors are extended, dimeric coiled-coil proteins characterized by a set of conserved motifs that promote the stabilization of the Dynein-Dynactin complex (Carter *et al.*, 2016; Olenick & Holzbaur, 2019; Reck-Peterson *et al.*, 2018). While conserved in their outline, adaptors respond to different stimuli in different subcellular locales and interact with different cargoes. Spindly is recognized as the kinetochore adaptor for DD (Chan *et al*, 2009; Gassmann, 2023; Griffis *et al*, 2007; Yamamoto *et al*, 2008). Its localization to kinetochores requires an interaction with the RZZ complex that is greatly enhanced by Spindly farnesylation (Gassmann, 2023; Kops & Gassmann, 2020).

In addition to promoting bi-orientation, the corona has also been shown to promote microtubule nucleation (Wu *et al*, 2023) and to contribute to the spindle assembly checkpoint (SAC), a pathway that delays anaphase onset until successful bi-orientation of all sister chromatid pairs (Fischer, 2022; Lara-Gonzalez *et al*, 2021). One of the main components of the SAC, the MAD1:MAD2 core complex, is also a constituent of the corona, where it also interacts with the CDK1:Cyclin B complex (Alfonso-Perez *et al*, 2019; Allan *et al*, 2020; Hoffman *et al*, 2001; Jackman *et al*, 2020). SAC signaling at each kinetochore subsides during the conversion of kinetochore attachments from the microtubule lattice to the microtubule end (Chakraborty *et al*, 2019; Kuhn & Dumont, 2017, 2019; Magidson *et al*, 2015; Sacristan *et al.*, 2018). Following this lateral to end-on conversion of kinetochore-microtubule attachment, which involves CENP-E (Chakraborty *et al.*, 2019), the corona is rapidly disassembled (shedding) (Basto *et al*, 2004; Hoffman *et al.*, 2001; Wojcik *et al*, 2001). Shedding is caused by activation of retrograde Dynein motility and promotes removal of corona components and their relocation to the spindle poles. This process also removes MAD1:MAD2 from the kinetochore, leading to suppression of SAC signaling (Ballister *et al*, 2014; Fava *et al*, 2011; Kuhn & Dumont, 2017, 2019; Maldonado & Kapoor, 2011).

Correct coordination of end-on attachment and checkpoint silencing through shedding is crucial for mitosis. The precise order of molecular events behind this coordination, however, remains unclear. Whether CENP-E is merely recruited to the kinetochore corona as an outermost terminal component or rather contributes to the stabilization of the corona and the localization and function of other proteins, most notably DD, is currently unknown. Thus, dissecting the interactions of CENP-E at the corona has become pressing. MAD1 has been proposed to act as a kinetochore receptor for CENP-E (Akera *et al*, 2015). Other studies, however, did not identify a role of MAD1 in CENP-E recruitment (Martin-Lluesma *et al*, 2002; Sharp-Baker & Chen, 2001). Binding to CENP-E has also been proposed to control activation of the kinase activity of BUBR1 in SAC control (Mao *et al*, 2003; Mao *et al*, 2005), but other studies have suggested BUBR1 is a pseudokinase devoid of catalytic activity (Breit *et al*, 2015; Suijkerbuijk *et al*, 2012). BUBR1 interacts directly with CENP-E and contributes to its kinetochore recruitment (Ciossani *et al*, 2018; Legal *et al*, 2020). We and others, however, observed that depletion of BUBR1 causes only modest reduction of CENP-E from prometaphase kinetochores (Akera *et al.*, 2015; Ciossani *et al.*, 2018; Lampson & Kapoor, 2005). Furthermore, unlike CENP-E, BUBR1 does not expand into the corona, suggesting that at least another prominent CENP-E receptor must be present in the corona.

A second set of pressing questions concerns the RZZS and its role in corona assembly. The recent realization that RZZS is the building block of the corona (Mosalaganti *et al.*, 2017; Pereira *et al.*, 2018; Raisch *et al.*, 2022; Sacristan *et al.*, 2018) raises questions on how the RZZS becomes recruited to kinetochores. For instance, it is unknown whether the requirements for kinetochore localization of the individual RZZS building blocks and of their polymer are the same. Furthermore, MPS1 kinase activity regulates corona expansion, but it is unclear whether it also regulates the interaction of RZZS with the kinetochore, as possibly implied by the reduced kinetochore levels of RZZS upon MPS1 inhibition (Rodriguez-Rodriguez *et al.*, 2018). Both the Ndc80C and the Knl1C have been implicated in RZZS recruitment (Caldas *et al*, 2015; Chan *et al.*, 2009; Lin *et al*, 2006; Pagliuca *et al*, 2009; Pereira *et al.*, 2018; Sundin *et al*, 2011; Varma *et al*, 2013), but other reports also identified these proteins as being at least partly dispensable (Pereira *et al.*, 2018; Rodriguez-Rodriguez *et al.*, 2018; Silio *et al*, 2015; Varma *et al.*, 2013).

Addressing how the kinetochore scaffold influences the corona is very challenging. Each of the two main regulatory branches of the KMN network, the Knl1C and the Ndc80C, individually recruits several regulatory proteins at the same time. These downstream regulators, however, mutually reinforce each other functionally and structurally. For instance, Ndc80C recruits MPS1 (Hiruma *et al*, 2015; Ji *et al*, 2015; Martin-Lluesma *et al.*, 2002; Stucke *et al*, 2004), but MPS1 phosphorylates Knl1C to promote recruitment of BUB1, which in turn recruits BUBR1 (Krenn *et al*, 2014; London *et al*, 2012; Meadows *et al*, 2011; Overlack *et al*, 2015; Primorac *et al*, 2013; Vleugel *et al*, 2013; Yamagishi *et al*, 2012). This complex connectivity, which is also expected to influence RZZS and CENP-E recruitment, is an unavoidable source of confusion when simple protein depletions are used as the main perturbation experiment. Therefore, dissecting this complexity necessitates the use of well-characterized separation-of-function mutants (whose identification is usually very laborious) and their re-introduction, possibly in the form of recombinant proteins or protein complexes delivered by protein electroporation (Alex *et al*, 2019; Polley *et al*, 2022) into cells depleted of the endogenous counterpart.

Here we break new ground in our dissection of the kinetochore corona. We demonstrate a direct interaction between CENP-E and the RZZS complex that makes them partly co-dependent for kinetochore localization and function. The interaction promotes kinetochore recruitment of DD, which is otherwise largely depleted in absence of CENP-E or in presence of a CENP-E mutant affecting RZZS binding. While we confirm the importance of BUBR1 in CENP-E recruitment, we find no clear evidence of a role of MAD1. Finally, we show that RZZS, in addition to CENP-E, requires KNL1, and Ndc80C to a minor extent, as kinetochore receptors, and that MPS1 is implicated not only in corona expansion, but also in the interaction of RZZS with the kinetochore. We discuss our results also in the context of the regulation of activation of opposing motor activities at different cellular locales.

## Results

### CENP-E interacts with BUBR1 and RZZS

BUBR1 depletion does not eliminate CENP-E from kinetochores (see Introduction). Because CENP-E localizes to the corona in prometaphase, we asked if perturbations of the corona also affected CENP-E localization. Individual RNAi based depletions of BUBR1 or Zwilch caused partial reductions of CENP-E in cells arrested in prometaphase with the spindle poison Nocodazole. Co-depletion, on the other hand, caused extensive reduction of CENP-E at kinetochores (Figure 1C-D). These results imply that both BUBR1 and the RZZS complex promote CENP-E localization. Residual CENP-E observed under conditions of co-depletion may reflect residual levels of BUBR1 and/or RZZ after the RNAi procedure (Figure 1-Supplement 1A-B), but we cannot exclude a weak interaction with a third receptor (see below).

Next, we tried to identify the kinetochore binding determinants of CENP-E. CENP-E features a C-terminal kinetochore-binding domain encompassing residues 2126-2476 (Chan *et al*, 1998) (Figure 1E). In our previous work, we demonstrated that a larger construct encompassing amino acids 2070-2701 (hereafter referred to as 2070C) recapitulates localization of full-length CENP-E and is sufficient for robust kinetochore localization in prometaphase (Ciossani *et al.*, 2018). Indeed, EGFP-CENP-E^2070C^ localized robustly to kinetochores in cells depleted of endogenous CENP-E (Figure 1F-G). This construct also elicited a robust mitotic checkpoint arrest as a consequence of widespread chromosome alignment defects (Figure 1-Supplement 1C-D).

Further dissection of CENP-E^2070C^ in N-terminal (2070-2365) and C-terminal (2366-2701, hereafter referred to as 2366C) fragments revealed that both decorated kinetochores in prometaphase-arrested cells depleted of endogenous CENP-E, even if at partially reduced levels relatively to CENP-E^2070C^ (Figure 1F-G). CENP-E^2070-2365^ localized to kinetochores without apparently extending into the corona. Conversely, CENP-E^2366C^ localized to corona crescents, suggesting that it might interact with RZZS or with another protein associated with the RZZS (Figure 1F-G). Thus, each fragment distinguishes unique determinants of kinetochore localization.

Depletion of BUBR1 prevented kinetochore recruitment of EGFP-CENP-E^2070-2365^ (the endogenous CENP-E was also depleted to prevent possible confounding effects of dimerization of endogenous and exogenous CENP-E) (Figure 1-Supplement 1E-G). Kinetochore localization of CENP-E^2070-2365^ may be mediated through an interaction with the pseudokinase domain of BUBR1 (Chan *et al.*, 1998; Ciossani *et al.*, 2018; Legal *et al.*, 2020; Yao *et al*, 2000). Indeed, analytical size-exclusion chromatography (SEC) confirmed a direct interaction of CENP-E^2070C^ or CENP-E^2070-2365^, but not CENP-E^2366C^, with the BUBR1 kinase domain (abbreviated as KD, Figure 1-Supplement 1H-I).

To identify the localization determinants of CENP-E^2366C^, we asked whether it interacts directly with RZZS. We polymerized mCherry-tagged RZZS *in vitro* to form minicircles, as described (Raisch *et al.*, 2022). We then imaged the minicircles by total internal reflection fluorescence (TIRF) microscopy while testing association with various recombinant EGFP-CENP-E constructs. Both EGFP-CENP-E^2070C^ and EGFP-CENP-E^2366C^ co-localized with the RZZS minicircles *in vitro*, while EGFP-CENP-E^2070-2365^ failed to do so (Figure 1H). The BUBR1 kinase domain also failed to compete with EGFP-CENP-E^2070C^ for co-localization on the RZZS minicircles (Figure 1I). These observations suggest that CENP-E^2366C^ contains a binding site for direct interaction with the RZZS complex.

### Determinants of CENP-E interactions with BUBR1 and RZZS

To dissect further the function of CENP-E^2070-2365^ and CENP-E^2366C^, we introduced mutations within conserved sequence stretches in these segments. Specifically, we targeted residues in the segments 2185-2195 and 2497-2507, residing respectively in the BUBR1-binding and RZZS-binding regions of CENP-E (Figure 2-Supplement 1A-B). Analytical SEC confirmed that charge-reversal mutations in the 2185-2195 motif abrogated binding of CENP-E^2070C^ to the pseudokinase domain of BUBR1 (Figure 2-Supplement 1C). Hereafter, we will refer to the mutant in the 2185-2195 region as the BUBR1^Mut^ of CENP-E. AlphaFold2 (Jumper *et al*, 2021) predicts this region to be juxtaposed, in the same 4- helix bundle, to residues within the 2310-2320 region whose mutation also abolishes BUBR1 binding (Legal *et al.*, 2020) (Figure 2-Supplement 1D).

**Figure 2.**
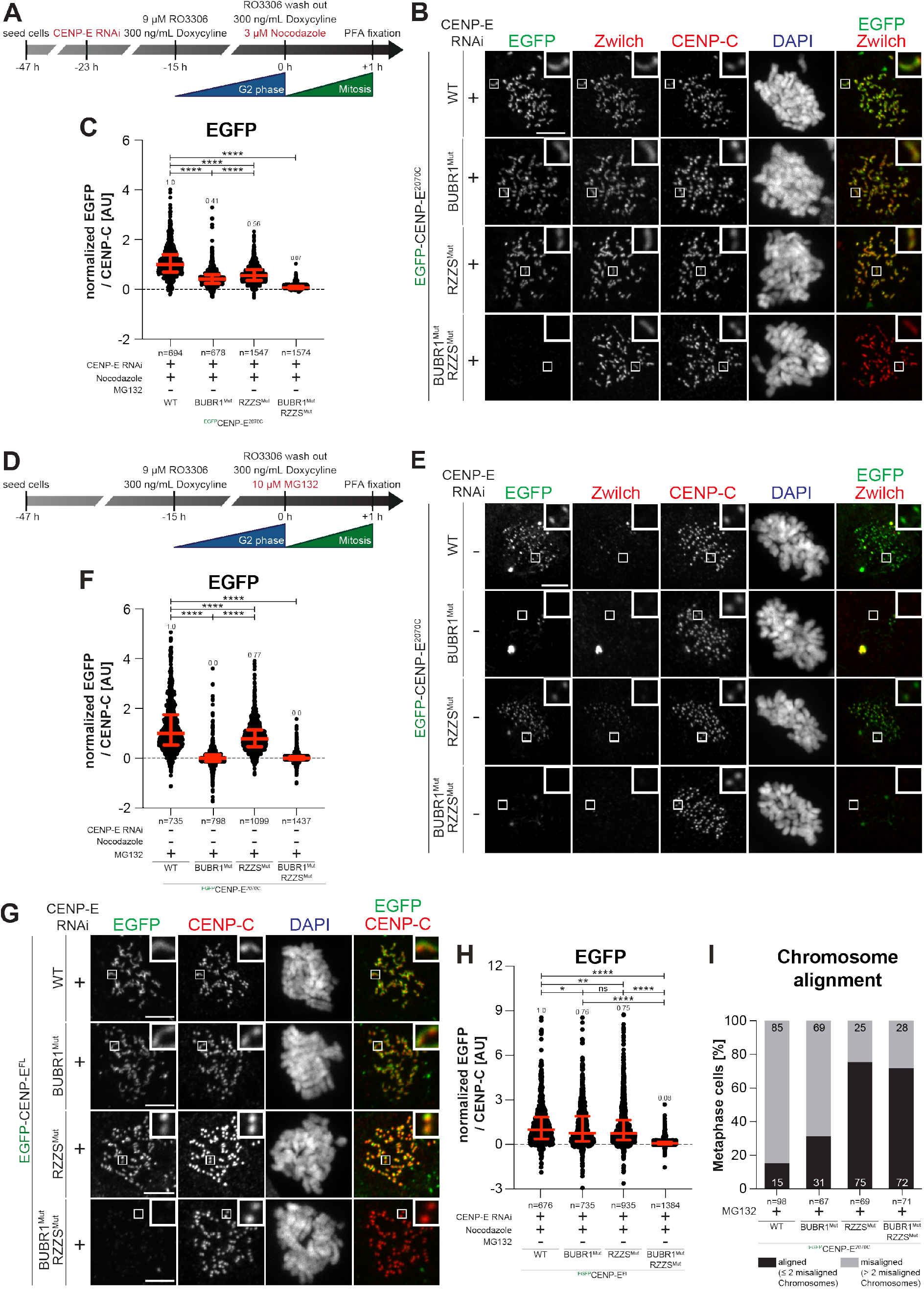
Separation of function mutants validate BUBR1 and RZZS as CENP-E partners. (**A**) Schematic representation of the experimental scheme used for panels B-C. (**B**) Representative images showing the localization of different ^EGFP^CENP-E^2070C^ mutants in stable DLD-1 cell lines arrested in prometaphase. CENP-E RNAi treatment was performed for 24 h with 60 nM siRNA. 8 h after RNAi treatment, protein expression was induced through addition of 300 ng/mL doxycycline and cells were synchronized in G2 phase with 9 μM RO3306 for 15 h and then released into mitosis. Subsequently, cells were immediately treated with 3.3 μM Nocodazole and 300 ng/mL doxycycline for an additional hour. CENP-C was used to visualize kinetochores and DAPI to stain DNA. Scale bar: 5 μm. (**C**) Quantification of EGFP levels at kinetochores of the experiment shown in panel B. n refers to individually measured kinetochores. (**D**) Schematic representation of the experimental scheme used for panels E-F. (**E**) Representative images showing the localization of different ^EGFP^CENP-E^2070C^ mutants in stable DLD-1 cell lines arrested in metaphase. 32 h after cells were seeded, protein expression was induced through addition of 300 ng/mL doxycycline and cells were synchronized in G2 phase with 9 μM RO3306 for 15 h and then released into mitosis. Subsequently, cells were immediately treated with 10 μM MG132 and 300 ng/mL doxycycline for two hours. CENP-C was used to visualize kinetochores and DAPI to stain DNA. Scale bar: 5 μm. (**F**) Quantification of EGFP levels at kinetochores of the experiment shown in panel E. n refers to individually measured kinetochores. (**G**) Representative images showing the localization of different ^EGFP^CENP-E^FL^ constructs in stable DLD-1 cell lines arrested in prometaphase. CENP-E RNAi treatment was performed for 24 h with 60 nM siRNA. 8 h after RNAi treatment protein expression was induced through addition of 10 ng/mL doxycycline and cells were synchronized in G2 phase with 9 μM RO3306 for 15 h and then released into mitosis. Subsequently, cells were immediately treated with 3.3 μM Nocodazole and 10 ng/mL doxycycline for an additional hour. CENP-C was used to visualize kinetochores and DAPI to stain DNA. Scale bar: 5 μm. (**H**) Quantification of EGFP levels at kinetochores of the experiment shown in panel G. n refers to individually measured kinetochores. (**I**) Chromosome alignment analysis of stable cell lines expressing different ^EGFP^CENP-E^2070C^ constructs. Before fixation, cells were synchronized in G2 phase with 9 μM RO3306 for 15 h and then released into mitosis. Subsequently, cells were immediately treated with 10 μM MG132 for two hours. n refers to the number of analyzed metaphase cells.

To test the effects of mutations in the 2497-2507 region, we formed RZZS filaments (Raisch *et al.*, 2022) and tested co-localization of EGFP-CENP-E^2070C^. A mutation of four conserved residues in the 2497-2507 region prevented interaction with RZZS in this assay (Figure 2-Supplement 1E). The binding defect was further substantiated in the TIRF assay, in which we compared binding of EGFP-CENP-E^2366C^ to RZZS minicircles. While the wild type EGFP-CENP-E^2366C^ construct bound minicircles, the mutant did not (Figure 2-Supplement 1F). Hereafter, we will therefore refer to the mutant in the 2497-2507 region as the RZZS^Mut^ of CENP-E.

We generated stable colorectal adenocarcinoma DLD-1 cell lines expressing EGFP-CENP-E^2070C^ and mutants at similar levels from an inducible promoter (Figure 2-Supplement 2A). EGFP-CENP-E^2070C^ carrying the BUBR1^Mut^ and RZZS^Mut^ localized at kinetochores in prometaphase-arrested cells, with the BUBR1^Mut^ decorating the corona, and the RZZS^Mut^ decorating kinetochores (Figure 2A-C). Conversely, a double mutant was unable to decorate kinetochores, implying that binding to at least one site is necessary for recruitment in prometaphase (Figure 2A-C). Essentially identical results were obtained when purified EGFP-CENP-E^2070C^ protein constructs were electroporated in cells (Figure 2-Supplement 2B-C). Thus, the BUBR1 and RZZS binding sites of CENP-E^2070C^ can promote robust recruitment of CENP-E even independently of each other.

Indeed, electroporated EGFP-CENP-E^2070-2365^ and EGFP-CENP-E^2366C^ constructs, respectively, localized to the kinetochore and the corona in cells depleted of endogenous CENP-E. Introduction of the BUBR1^Mut^ in EGFP-CENP-E^2070-2365^ or of the RZZS^Mut^ in EGFP-CENP-E^2366C^ prevented their kinetochore recruitment (Figure 2-Supplement 2D-G).

As previously shown (Gassmann *et al*, 2010) the kinetochore levels of BUBR1 and Zwilch decrease upon bi-orientation; however, while the kinetochore levels of BUBR1 remained comparatively high, Zwilch was only present at trace levels (quantifications shown in Figure 2-Supplement 2H-I). Accordingly, CENP-E was partially retained at kinetochores after corona shedding and achievement of bi-orientation (Ciossani *et al.*, 2018). Thus, residual CENP-E on metaphase kinetochores may localize exclusively through BUBR1. Indeed, when localization experiments with BUBR1^Mut^ and RZZS^Mut^ of EGFP-CENP-E^2070C^ were performed in metaphase-arrested cells (through addition of the proteasome inhibitor MG132), the RZZS^Mut^ decorated kinetochores whereas the BUBR1^Mut^ was depleted (Figure 2D-F). Also in this case, the double mutant failed to decorate kinetochores. Thus, after corona shedding, BUBR1 is the only residual CENP-E receptor at the kinetochore, so that only the RZZS^Mut^ can retain kinetochore localization at metaphase. Localization patterns that were essentially identical to those of EGFP-CENP-E^2070C^ were also obtained with full-length CENP-E and its BUBR1^Mut^ and RZZS^Mut^ mutants expressed from inducible DLD-1 cell lines (Figure 2G-H).

In line with the localization experiments, expression of CENP-E^2070C^ had a severe dominant effect on chromosome alignment, likely because the exogenous protein displaces endogenous CENP-E (Figure 2I). Conversely, the BUBR1^Mut^ of CENP-E^2070C^ had less prominent effects, and the RZZS^Mut^ or the double mutant had almost no effect on chromosome alignment, suggesting that they have very limited capacity to displace endogenous CENP-E. These observations suggest that simultaneous interactions with RZZS and BUBR1 stabilize CENP-E.

### MPS1 inhibition exposes a role of CENP-E in RZZS localization

CENP-E co-localizes with the fibrous corona in prometaphase and is partially removed from the kinetochore upon end-on attachment (Ciossani *et al.*, 2018; Cooke *et al*, 1997; Yao *et al*, 1997). Initially, we asked if CENP-E contributes to kinetochore recruitment or retention of RZZS in prometaphase-arrested cells. To this end, we used an hTERT-immortalized retinal pigment epithelial-1 (hTERT-RPE-1) cell line in which both endogenous CENP-E alleles were C-terminally tagged with an auxin-inducible degron (AID) and a 3xFLAG tag (Owa & Dynlacht, 2021). In untreated control cells, CENP-E adopted the characteristic crescent shape of kinetochore coronas. Addition of the auxin derivative indole acetic acid (IAA) caused rapid degradation of CENP-E to undetectable levels within 20 minutes (Figure 3B) (Owa & Dynlacht, 2021). We then monitored RZZS levels in mitotic cells where CENP-E had been degraded either before or after mitotic entry (respectively indicated as T_-30_ and T_+30_ in Figure 3A, C-F). Irrespective of the degradation protocol used, these experiments did not reveal large changes in RZZS kinetochore levels upon degradation of CENP-E (Figure 3E-F). Similar results were obtained after RNAi-mediated depletion of CENP-E in HeLa cells (Figure 3-Supplement 1A-D). Thus, the kinetochore corona remains stable without CENP-E.

**Figure 3.**
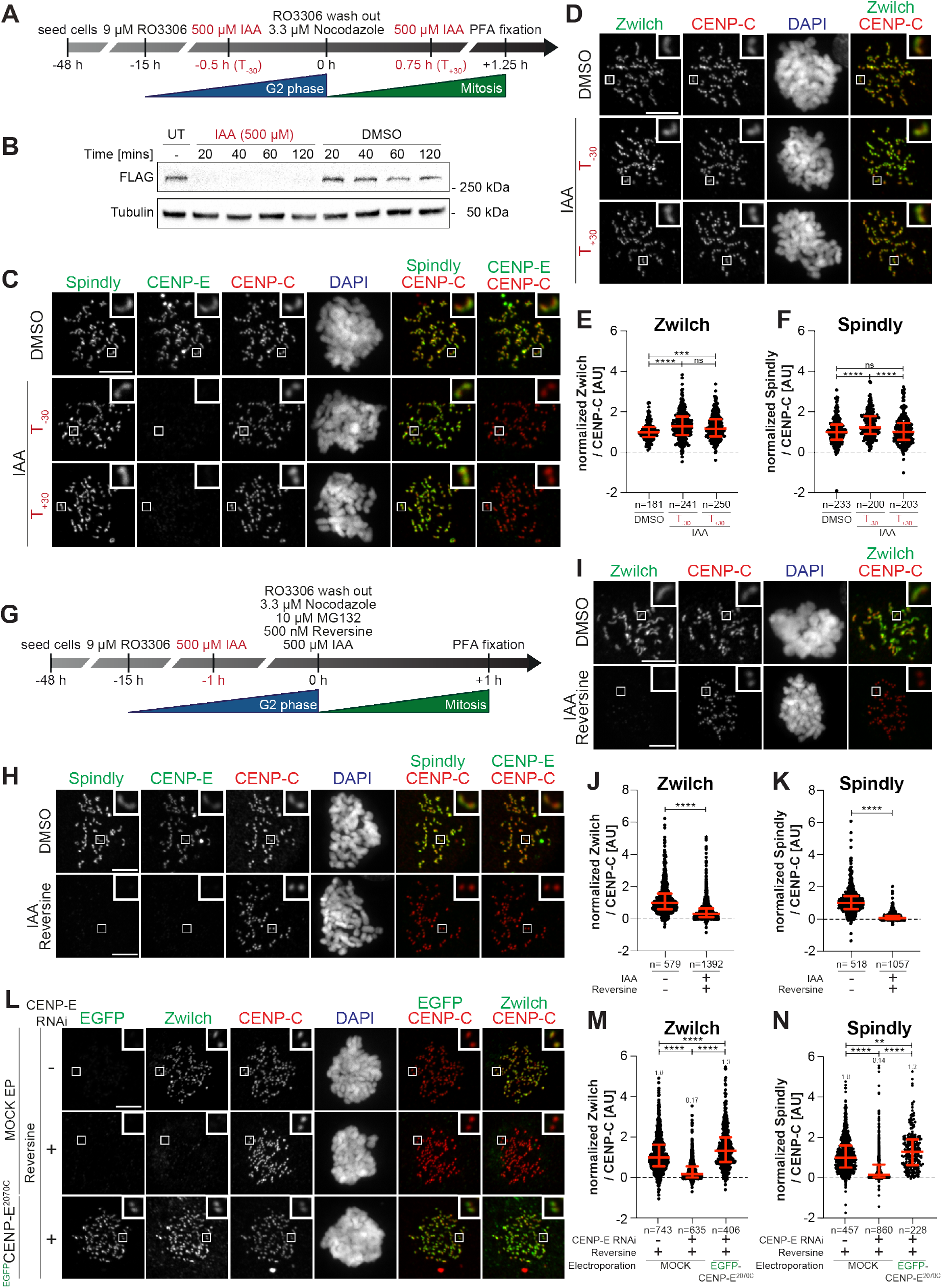
Combining CENP-E depletion and MPS1 inhibition dissolves the corona. (**A**) Schematic representation of the cell synchronization protocols for the experiment in panels C-F. (**B**) Immunoblot of mitotic RPE-1 cells showing the levels of endogenous CENP-E^AID_3xFLAG^ treated with 500 µM IAA for the indicated duration. 50 μg of cleared lysate was used for each condition, and Tubulin is shown as a loading control. (**C-D**) Representative images showing the effect of degrading the endogenous CENP-E in RPE-1 cells before or after mitotic entry, as shown in panel A. Before fixation, cells were synchronized in G2 phase with 9 µM RO3306, released into mitosis and immediately treated with 3.3 µM Nocodazole. Cells were treated either 30 mins before or 45 mins after mitotic entry with 500 µM IAA. CENP-C was used to visualize kinetochores and DAPI to stain DNA. Scale bar: 5 μm. (**E-F**) Quantification of kinetochore levels of Zwilch and Spindly in cells depleted of the endogenous CENP-E as shown in panels C-D. n refers to individually measured kinetochores. (**G**) Schematic representation of the cell synchronization protocol for the experiment shown in panels H-K. (**H-I**) Representative images showing the effects of degrading the endogenous CENP-E in RPE-1 cells before mitotic entry, as shown in panel G. Before fixation, cells were synchronized in G2 phase with 9 µM RO3306, released into mitosis and immediately treated with 3.3 µM Nocodazole, 10 µM MG132 and 500 nM Reversine. Scale bar: 5 μm. (**J-K**) Quantification of kinetochore levels of Zwilch and CENP-E in cells depleted of the endogenous CENP-E and treated with Reversine as shown in panels H-I. n refers to individually measured kinetochores. (**L**) Representative images showing the localization of Zwilch in prometaphase after depletion of CENP-E with 60 nM siRNA. 13 h after RNAi treatment, cells were electroporated with electroporation buffer or ^EGFP^CENP-E^2070C^. Following an 8 h recovery, cells were synchronized in G2 phase with 9 μM RO3306 for 15 h and then released into mitosis. Subsequently, cells were immediately treated with 3.3 μM Nocodazole, 10 μM MG132 and, where indicated, with 500 nM Reversine, for an additional hour. CENP-C was used to visualize kinetochores and DAPI to stain DNA. Scale bar: 5 μm. (**M-N**) Quantification of Zwilch and Spindly levels at kinetochores of the experiment shown in panel L.

MPS1 kinase phosphorylates ROD on Thr13 and Ser15, and MPS1 inhibition prevents corona assembly while causing only relatively minor reductions of the kinetochore levels of the RZZ complex (Raisch *et al.*, 2022; Rodriguez-Rodriguez *et al.*, 2018; Sacristan *et al.*, 2018). While our observations indicate that CENP-E is not required for RZZS recruitment when corona expansion proceeds normally, CENP-E may contribute to RZZS recruitment before corona expansion. To assess this, we examined RZZS levels after depletion of CENP-E either in presence of MPS1 kinase activity, or after its inhibition to prevent corona expansion. In agreement with previous reports (Raisch *et al.*, 2022; Rodriguez-Rodriguez *et al.*, 2018), the specific MPS1 small-molecule inhibitor Reversine (Santaguida *et al*, 2010) slightly reduced the kinetochore levels of RZZS and prevented corona expansion (Figure 3-Supplement 1A-D). Next, we synchronized RPE-1 CENP-E^AID^ cells in G2 phase, through addition of the small-molecule RO3306, and added IAA one hour before release into mitosis to ensure complete degradation of CENP-E (Figure 3G). Concomitant depletion of CENP-E and inhibition of MPS1 caused RZZS to disappear from the kinetochore (Figure 3H-K). RNAi-mediated depletion of CENP-E combined with Reversine treatment resulted in essentially identical observations (Figure 3-Supplement 1A-D).

Recombinant EGFP-CENP-E^2070C^ protein electroporated in HeLa cells depleted of endogenous CENP-E and treated with Reversine was sufficient to restore robust RZZS localization (Figure 3L-N). EGFP-CENP-E^2366C^, on the other hand, localized normally in absence of endogenous CENP-E, but further inhibition of MPS1 prevented its kinetochore recruitment and caused a very strong reduction in the kinetochore levels of RZZS (Figure 3-Supplement 1E-G). These observations are consistent with the idea that kinetochore recruitment of RZZS depends on CENP-E when MPS1 activity is inhibited. They further demonstrate that to be effective in maintaining RZZS at the kinetochore, CENP-E may need to interact at the same time with RZZS and another receptor, most likely BUBR1, that CENP-E^2366C^ does not recognize. Collectively, these results indicate that CENP-E is dispensable for holding the RZZS onto kinetochores after corona expansion, but is essential for RZZS recruitment when corona assembly is inhibited with an MPS1 inhibitor, a previously unappreciated co-dependence of RZZS and CENP-E for kinetochore recruitment.

IAA-mediated destruction of CENP-E in hTERT-RPE-1 cells did not cause elimination of MAD1, which remained strongly bound to kinetochores (Figure 3-Supplement 1H-I). Treatment with Reversine, on the other hand, caused a strong decrease in MAD1 kinetochore levels. No corresponding decrease in the levels of CENP-E was observed (Figure 3-Supplement 1J-L), however, an observation that seems inconsistent with the proposed role of MAD1 as a CENP-E receptor (Akera *et al.*, 2015).

### CENP-E contributes to kinetochore recruitment of Dynein-Dynactin

While CENP-E becomes dispensable for robust corona expansion after MPS1-mediated phosphorylation of RZZS, it may continue to interact with, and influence the function of, the RZZS even after corona expansion. As discussed in the Introduction, a primary function of RZZS is to bind and activate Dynein-Dynactin (DD). DD adaptors have been previously proposed to be a fulcrum for co-regulation of DD and kinesin activity (see Discussion). Thus, we decided to investigate if CENP-E influenced DD activity at the kinetochore. Supporting this hypothesis, RNAi-mediated depletion of CENP-E caused a very substantial reduction of p150^glued^ (a subunit of Dynactin, hereafter simply referred to as p150) at kinetochores in prometaphase arrested cells (Figure 4A-D). Normal levels of p150 were restored after electroporation of recombinant CENP-E^2070C^ but not of the RZZS^Mut^ mutant, indicating that binding of CENP-E to the RZZS complex is required for accumulation of DD (Figure 4A-D). Thus, in addition to the RZZS complex, CENP-E also contributes substantially to DD recruitment, and possibly activation, at the kinetochore. Of note, these changes in the levels of DD did not affect the levels of Spindly, which remained largely unaltered (quantified in Figure 4D), implying that CENP-E does not contribute to DD recruitment by controlling the levels of Spindly.

**Figure 4.**
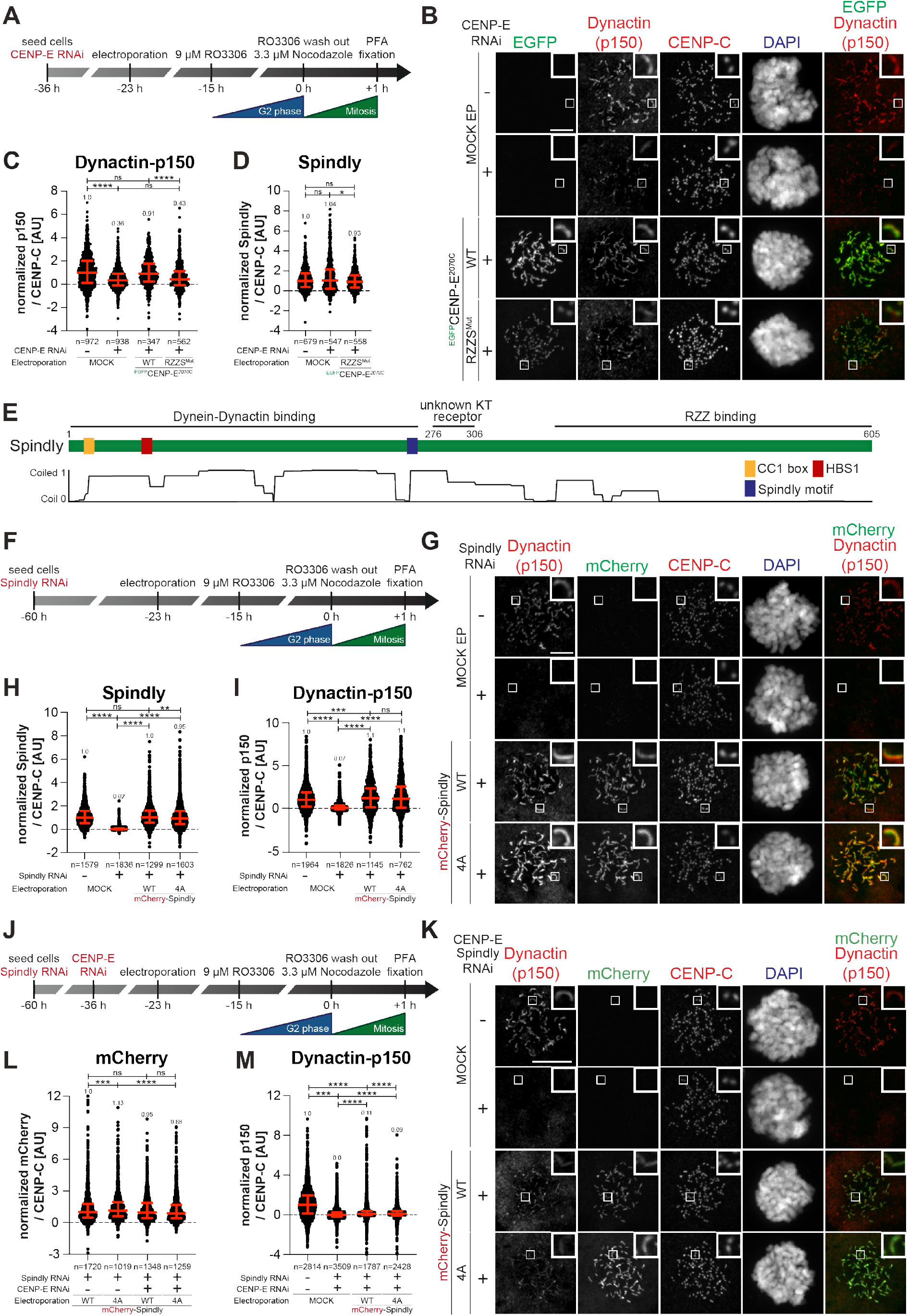
CENP-E is important for Dynein-Dynactin kinetochore recruitment. (**A**) Schematic representation of the cell synchronization protocols for the experiment in panels B-D. (**B**) Representative images showing the localization of Dynactin monitored through the p150^glued^subunit after depletion of CENP-E with 60 nM siRNA. 13 h after RNAi treatment cells were electroporated with electroporation buffer or recombinant ^EGFP^CENP-E^2070C^ constructs as indicated. Following an 8 h recovery, cells were synchronized in G2 phase with 9 μM RO3306 for 15 h and then released into mitosis. Subsequently, cells were immediately treated with 3.3 μM Nocodazole for an additional hour. CENP-C was used to visualize kinetochores and DAPI to stain DNA. Scale bar: 5 μm. (**C-D**) Quantification of Dynactin-p150^glued^ and Spindly levels at kinetochores of the experiment shown in panel B. n refers to individually measured kinetochores. (**E**) Schematic representation of the organization of Spindly and relevant coiled-coil prediction. (**F**) Schematic representation of the cell synchronization protocols for the experiment in panels G-I. (**G**) Representative images showing the localization of Dynactin monitored through the p150^glued^ subunit after depletion of Spindly with 50 nM siRNA. 37 h after RNAi treatment, cells were electroporated with electroporation buffer or recombinant ^mCh^Spindly constructs as indicated. Following an 8 h recovery, cells were synchronized in G2 phase with 9 μM RO3306 for 15 h and then released into mitosis. Subsequently, cells were immediately treated with 3.3 μM Nocodazole for an additional hour. CENP-C was used to visualize kinetochores and DAPI to stain DNA. Scale bar: 5 μm. (**H-I**) Quantification of Spindly and Dynactin-p150^glued^ levels at kinetochores of the experiment shown in panel G. n refers to individually measured kinetochores. (**J**) Schematic representation of the cell synchronization protocols for the experiment in panels K-M. (**K**) Representative images showing the localization of Dynactin monitored through the p150^glued^ subunit after depletion of CENP-E with 60 nM siRNA and Spindly with 50 nM siRNA. 13 h after CENP-E RNAi treatment cells were electroporated with electroporation buffer or recombinant ^mCh^Spindly constructs as indicated. Following an 8 h recovery, cells were synchronized in G2 phase with 9 μM RO3306 for 15 h and then released into mitosis. Subsequently, cells were immediately treated with 3.3 μM Nocodazole for an additional hour. CENP-C was used to visualize kinetochores and DAPI to stain DNA. Scale bar: 5 μm. (**L-M**) Quantification of mCherry and Dynactin-p150^glued^ levels at kinetochores of the experiment shown in panel K. n refers to individually measured kinetochores.

Spindly contains various sequence elements implicated in its function and regulation as a DD adaptor (Figure 4E). In previous work, we reported that Spindly may exist in an autoinhibited conformation (Figure 4-Supplement 1A) and proposed that an unknown kinetochore trigger promotes a conformational change required for efficient binding of Spindly to DD at the kinetochore (d’Amico *et al*, 2022). The Spindly motif, identified in Spindly and other DD adaptors (Figure 4E), promotes an interaction with the pointed-end (PE) subcomplex of Dynactin (Gama *et al*, 2017; Gassmann *et al.*, 2010). In agreement with our model that Spindly is natively autoinhibited, mCherry-Spindly did not bind PE in SEC experiments (Figure 4-Supplement 1B).

In our previous work, we also described mutants of Spindly that overcome autoinhibition and bind PE (d’Amico *et al.*, 2022). These mutants, however, were not recruited effectively to kinetochores, presumably because the mutated residues (in the 276-306 region of Spindly) also affected the interaction of Spindly with an unknown kinetochore receptor (Barisic *et al*, 2010; d’Amico *et al.*, 2022). We therefore scanned additional mutants to identify a separation-of-function mutant that would relieve Spindly autoinhibition without affecting kinetochore recruitment. Contrary to Spindly^WT^, a Spindly^4A^ mutant (described in Figure 4-Supplement 1A) interacted with the PE complex in a SEC experiment (Figure 4-Supplement 1B-C), suggesting autoinhibition has been relieved. Importantly, in cells depleted of endogenous Spindly, electroporated mCherry-Spindly^4A^ localized to the corona indistinguishably from its wild type counterpart and recruited DD (Figure 4F-I).

These observations suggest that we might have obtained an open mutant of Spindly that can also be efficiently recruited to the kinetochore, thus outperforming our previously described mutants (d’Amico *et al.*, 2022). We speculated that upon binding of Spindly at the kinetochore, CENP-E may trigger the conformational change that relieves Spindly auto-inhibition. In SEC experiments, Spindly^4A^, but not Spindly^WT^, interacted with CENP-E, albeit weakly (Figure 4-Supplement 1D-E). Being at least partially relieved of autoinhibition, Spindly^4A^ may be expected to rescue the defect in DD recruitment caused by depletion of CENP-E. Contrary to this hypothesis, however, Spindly^4A^, even if normally recruited to kinetochores in absence of CENP-E, did not rescue the defect on DD recruitment caused by CENP-E depletion (Figure 4J-M). Thus, Spindly^4A^ is insufficient to rescue the effects on Dynactin recruitment caused by depletion of CENP-E. Another open Spindly construct, Spindly^33-605^ (lacking the first 32 residues containing the Spindly CC1 box, also required for autoinhibition) (d’Amico *et al.*, 2022) also localized normally to kinetochores and rescued Dynactin levels upon depletion of endogenous Spindly. Yet, like Spindly^4A^, this mutant was likewise unable to recruit Dynactin in absence of CENP-E (quantified in Figure 4-Supplement 1F-G). Collectively, these results suggest either that Spindly^4A^ and Spindly^33-605^ are not fully “open” or that they are, but CENP-E is additionally required for DD recruitment.

As depletion of CENP-E led to a very significant but incomplete depletion of Dynactin, we asked if the residual Dynactin was recruited through CENP-F, which has been shown to participate in the recruitment of DD to the kinetochore and corona compaction (Gassmann, 2023; Mitevska *et al*, 2023; Vergnolle & Taylor, 2007). While CENP-F depletion had insignificant effects on the kinetochore levels of Dynactin, its combination with CENP-E depletion led to an almost complete depletion of kinetochore Dynactin (Figure 4-Supplement 1H-K), with the residual Dynactin signal probably reflecting recruitment by Spindly. Thus, recruitment of DD to the kinetochore may reflect concomitant interactions with RZZS, CENP-E, and CENP-F.

### R^EE^ZZ bypasses requirements for CENP-E and MPS1 kinase activity

RZZS depends on CENP-E for its kinetochore localization when MPS1 is inhibited. We reasoned that this requirement for CENP-E to recruit RZZS in absence of MPS1 kinase activity would be bypassed if MPS1 phosphorylation triggering of corona expansion could be mimicked. Phosphorylation of T13 and S15 on ROD by MPS1 is a prerequisite for corona expansion and RZZS polymerization (Raisch *et al.*, 2022; Rodriguez-Rodriguez *et al.*, 2018). In filamentation assays *in vitro*, mutation of these two residues to glutamic acid (E) allows RZZS polymerization in the absence of MPS1, while mutation to alanine (A) prevents polymerization even in presence of MPS1 kinase activity (Raisch *et al.*, 2022). Thus, we performed experiments to assess if a mutant RZZ carrying T13E and S15E mutations on ROD (hereafter referred to as R^EE^ZZ) bypasses a requirement for MPS1 activity for corona polymerization in cells. Contrary to electroporated wild type mCherry-RZZ, which was recruited to kinetochores but did not expand a corona in the presence of Reversine, electroporated mCherry-R^EE^ZZ expanded in a crescent shape even after inhibition of MPS1 (Figure 5A; expansion of the corona indicates that electroporated RZZ interacts with endogenous Spindly). Importantly, electroporated mCherry-R^EE^ZZ succeeded in assembling the corona even in Reversine treated cells additionally depleted of CENP-E and Zwilch by RNAi (Figure 5C-D). This observation provides an unequivocal demonstration that corona assembly harnesses a kinetochore receptor distinct from CENP-E. R^EE^ZZ was removed in a timely manner upon bi-orientation (Figure 5-Supplement 1A-D), suggesting that reversal of MPS1 phosphorylation of ROD may not be strictly required for corona disassembly.

**Figure 5.**
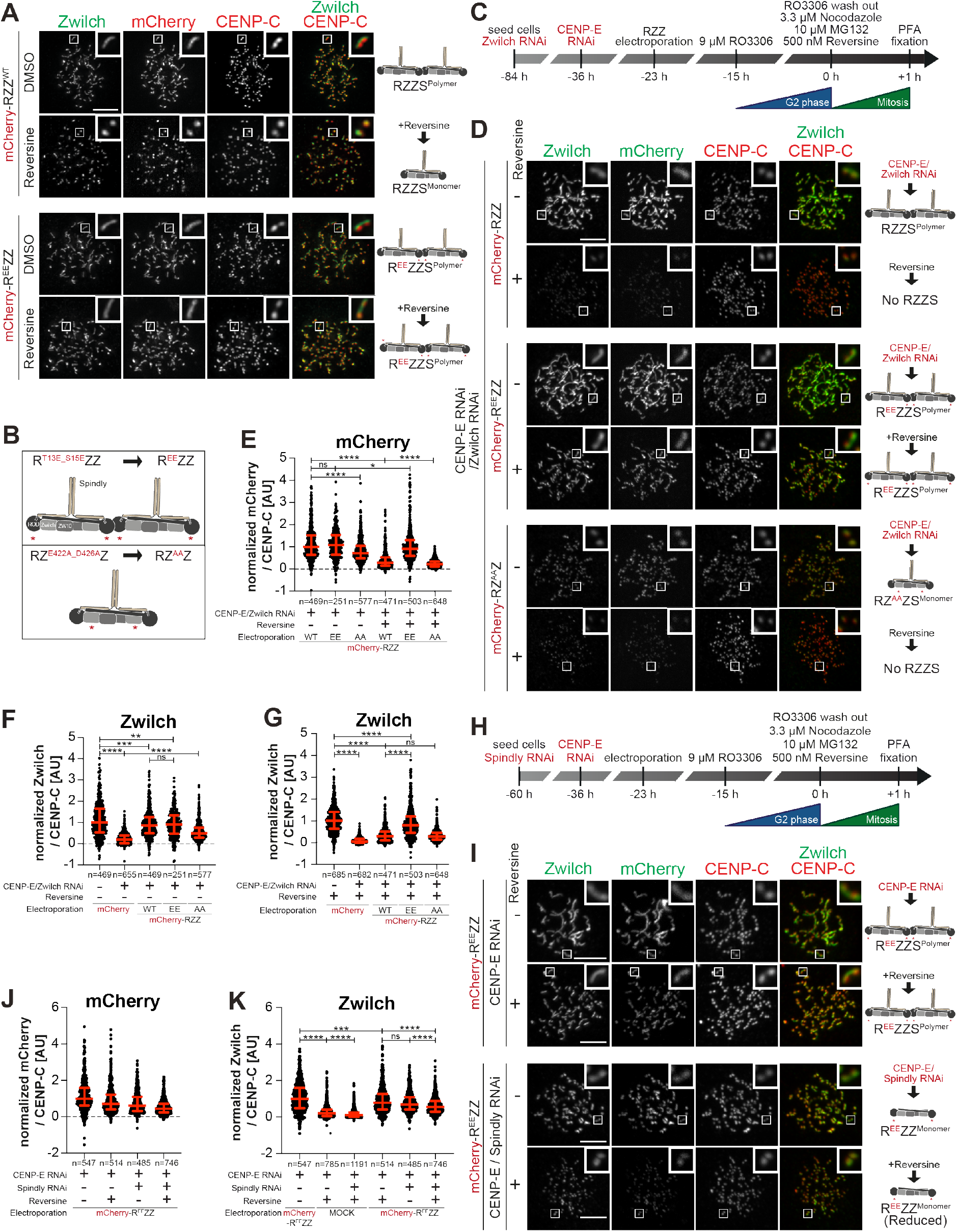
MPS1 contributes to RZZS recruitment in addition to corona expansion. (**A**) Representative images showing cells electroporated with the indicated ^mCh^RZZ constructs. Before fixation, cells were synchronized in G2 phase with 9 μM RO3306 for 15 h and then released into mitosis. Subsequently, cells were immediately treated with 3.3 μM Nocodazole, 10 μM MG132 and, where indicated, with 500 nM Reversine, for an additional hour. CENP-C was used to visualize kinetochores and DAPI to stain DNA. Scale bar: 5 μm. (**B**) Schematic of the ^mCh^RZZ constructs utilized in panel D. (**C**) Schematic of the cell synchronization and imaging experiment shown in panels D-G. (**D**) Representative images showing the localization of Zwilch in prometaphase after depletion of CENP-E with 60 nM siRNA and Zwilch with 100 nM siRNA as shown schematically in panel C. 13 h after CENP-E RNAi treatment cells were electroporated with mCherry or different ^mCh^RZZ constructs as indicated. Following an 8 h recovery, cells were synchronized in G2 phase with 9 μM RO3306 for 15 h and then released into mitosis. Subsequently, cells were immediately treated with 3.3 μM Nocodazole, 10 μM MG132 and, where indicated, with 500 nM Reversine, for an additional hour. CENP-C was used to visualize kinetochores and DAPI to stain DNA. Scale bar: 5 μm. (**E-G**) Quantification of mCherry and Zwilch levels at kinetochores of the experiment shown in panel D. n refers to individually measured kinetochores. (**H**) Schematic of the cell synchronization and imaging experiment shown in panels I-K. (**I**) Representative images showing the localization of Zwilch in prometaphase after depletion of CENP-E with 60 nM siRNA and Spindly with 50 nM siRNA as shown schematically in panel H. 13 h after CENP-E RNAi treatment cells were electroporated with electroporation buffer or ^mCh^R^EE^ZZ. Following an 8 h recovery, cells were synchronized in G2 phase with 9 μM RO3306 for 15 h and then released into mitosis. Subsequently, cells were immediately treated with 3.3 μM Nocodazole, 10 μM MG132 and, where indicated, with 500 nM Reversine, for an additional hour. CENP-C was used to visualize kinetochores and DAPI to stain DNA. Scale bar: 5 μm. (**J-K**) Quantification of mCherry and Zwilch levels at kinetochores of the experiment shown in panel I. n refers to individually measured kinetochores.

### MPS1 promotes kinetochore recruitment of RZZS

Besides preventing corona expansion, MPS1 inhibition causes a reduction in the kinetochore levels of RZZS (Figure 3-Supplement 1A-D). Whether this reduction reflects a second role of MPS1 (in addition to corona expansion) in promoting kinetochore localization of RZZS is unknown. To investigate this, we tried to block corona expansion by means other than MPS1 inhibition. In humans and nematodes corona expansion requires a negatively charged cluster on Zwilch that contains two conserved residues, Glu422 and Asp426 (Figure 5B). Mutation of these residues to alanine prevents corona expansion (Gama *et al.*, 2017; Pereira *et al.*, 2018). Furthermore, RZ^E422A/D426A^Z (hereafter referred to as RZ^AA^Z) bound Spindly but was unable to form polymers *in vitro* (Raisch *et al.*, 2022). In line with these previous observations, RZ^AA^Z electroporated in cells depleted of endogenous Zwilch decorated kinetochores, albeit at slightly reduced levels, but was unable to promote corona expansion (Figure 5-Supplement 1E-F). We then applied the protocol described in Figure 5C to compare the localization of mCherry-RZZ^WT^, mCherry-R^EE^ZZ, and mCherry-RZ^AA^Z in presence or absence of Reversine in cells depleted of endogenous CENP-E and RZZ. Contrary to mCherry-R^EE^ZZ, MPS1 inhibition entirely prevented RZ^AA^Z from being recruited to the kinetochore (Figure 5D-G). Thus, RZ^AA^Z decouples the effects of inhibiting MPS1 from those resulting from inhibition of corona expansion. Its behavior suggests that MPS1, in addition to corona expansion, additionally promotes kinetochore recruitment of RZZ. Robust kinetochore recruitment of the phosphomimetic mCherry-R^EE^ZZ mutant despite MPS1 inhibition also suggests that this additional function of MPS1 is dispensable if corona expansion can proceed.

To confirm this conclusion, we turned to two additional conditions known to prevent corona expansion, the mutation of T13 and S15 to non-phosphorylatable alanine residues and the depletion of Spindly (Raisch *et al.*, 2022; Rodriguez-Rodriguez *et al.*, 2018). Electroporated R^T13A/S15A^ZZ (abbreviated as R^AA^ZZ and not to be confused with RZ^AA^Z) decorated kinetochores but did not expand the corona in cells depleted of CENP-E and of endogenous RZZ. Confirming our model, inhibition of MPS1 caused an essentially complete depletion of R^AA^ZZ from kinetochores (Figure 5-Supplement 1G-I). Depletion of Spindly in cells also depleted of CENP-E was also compatible with robust RZZ recruitment, but corona expansion was inhibited. Further inhibition of MPS1 led to a complete depletion of endogenous RZZ from the kinetochore (Figure 5-Supplement 2A-B). Even in cells harboring electroporated R^EE^ZZ, depletion of Spindly and CENP-E prevented corona expansion. Further addition of Reversine caused a strong reduction in the kinetochore levels of R^EE^ZZ. Thus, in absence of corona polymerization, even R^EE^ZZ becomes sensitive to MPS1 inhibition for kinetochore localization (Figure 5H-K).

### Kinetochore recruitment of RZZS

Collectively, these results indicate that MPS1, in addition to promoting corona expansion, also promotes the interaction of RZZS with one or more kinetochore receptors other than CENP-E. To shed light on the mechanism of kinetochore targeting of RZZS, we initially assessed the role of the outer kinetochore in RZZS recruitment. Individual RNAi-based depletions of the Ndc80C and KNL1 caused large reductions of CENP-E and RZZ at kinetochores. Co-depletion of KNL1 and Ndc80C, on the other hand, led to the complete disappearance of RZZS and CENP-E (Figure 6-Supplement 1A-C). Thus, both KMN subcomplexes contribute to the recruitment of CENP-E and of RZZS, as discussed in the Introduction.

**Figure 6.**
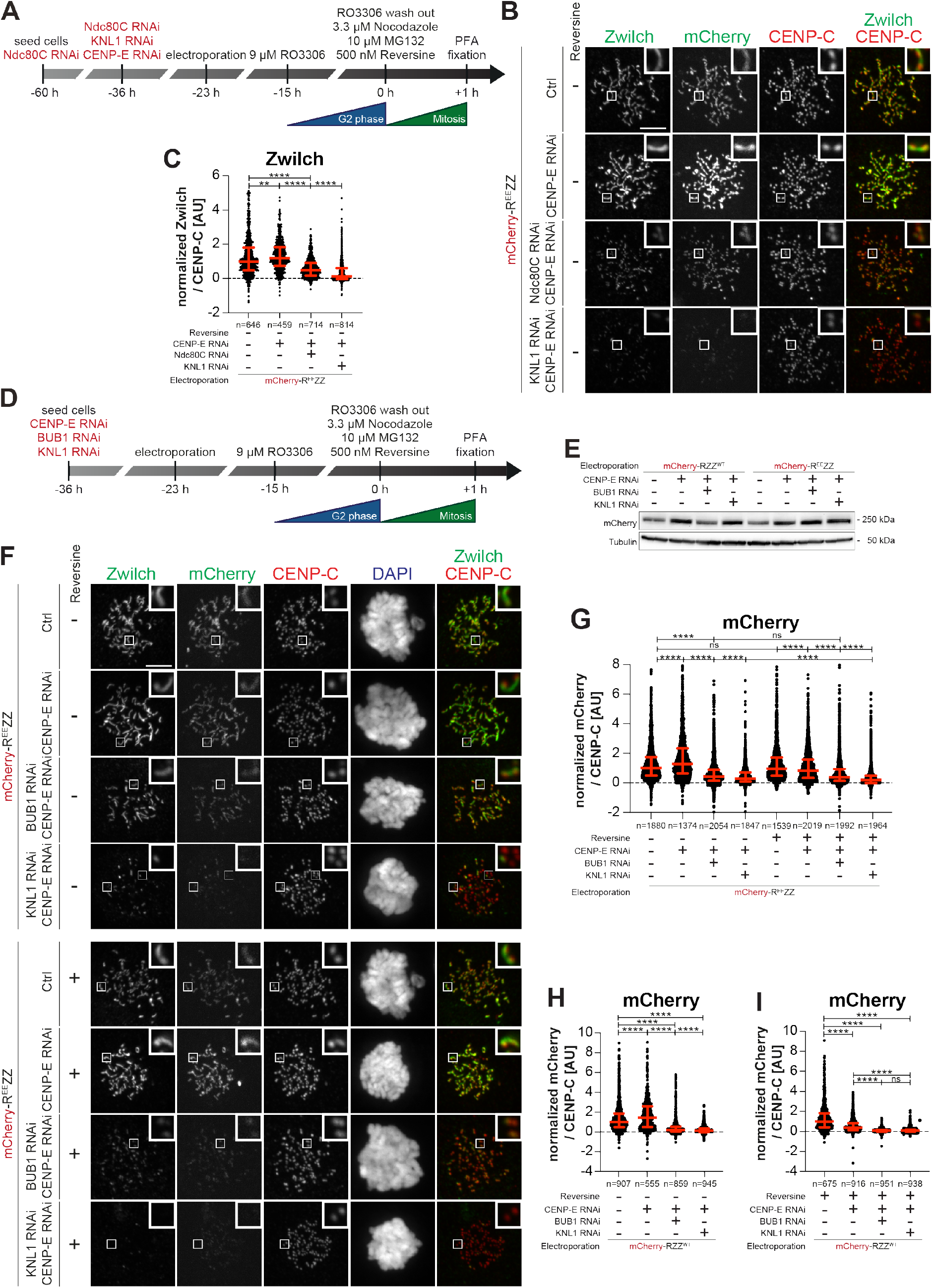
BUB1 and KNL1 are the main RZZS receptors. (**A**) Schematic of the cell synchronization and imaging experiment shown in panels B-C. (**B**) Representative images showing cells electroporated with ^mCh^R^EE^ZZ. Ndc80C RNAi treatment was performed with two transfections of 20 nM siRNA directed against HEC1/NDC80, SPC24, and SPC25 subunit. KNL1 and CENP-E RNAi were performed with 60 nM siRNA. 13 h after the second siNdc80C RNAi treatment and CENP-E RNAi or KNL1 RNAi treatment cells were electroporated with ^mCh^R^EE^ZZ. Following an 8 h recovery, cells were synchronized in G2 phase with 9 μM RO3306 for 15 h and then released into mitosis. Subsequently, cells were immediately treated with 3.3 μM Nocodazole and 10 μM MG132 for an additional hour. CENP-C was used to visualize kinetochores and DAPI to stain DNA. Scale bar: 5 μm. (**C**) Quantification of Zwilch levels at kinetochores of the experiment shown in panel B. n refers to individually measured kinetochores. (**D**) Schematic of the cell synchronization and imaging experiment shown in panel E-I. (**E**) Immunoblot of mitotic HeLa cells treated as shown in panel D and probed with the indicated antibodies. 50 μg of cleared lysate was used for each condition, and Tubulin is shown as a loading control. (**F**) Representative images showing cells electroporated with ^mCh^R^EE^ZZ as shown in panel D. BUB1 RNAi was performed with 50 nM, KNL1 and CENP-E RNAi were performed with 60 nM siRNA. 13 h after BUB1, KNL1, CENP-E RNAi treatment, cells were electroporated with ^mCh^R^EE^ZZ. Following an 8 h recovery, cells were synchronized in G2 phase with 9 μM RO3306 for 15 h and then released into mitosis. Subsequently, cells were immediately treated with 3.3 μM Nocodazole and 10 μM MG132 and, where indicated, with 500 nM Reversine, for an additional hour. CENP-C was used to visualize kinetochores and DAPI to stain DNA. Scale bar: 5 μm. (**G-I**) Quantification of mCherry levels at kinetochores of the experiment shown in panel F and for an equivalent experiment carried out with mCherry-RZZ^WT^ instead of mCherry-R^EE^ZZ. n refers to individually measured kinetochores.

These results do not necessarily imply a direct role of Ndc80C and KNL1 in RZZS recruitment, however. First, Ndc80C and Knl1C sustain each other reciprocally at the kinetochore, with a strong stabilization of Knl1C by Ndc80C and a less pronounced stabilization of Ndc80C by Knl1C (quantified in Figure 6-Supplement 1D-E). Furthermore, Ndc80C is required for kinetochore recruitment of MPS1 kinase (Hiruma *et al.*, 2015; Ji *et al.*, 2015; Martin-Lluesma *et al.*, 2002; Stucke *et al.*, 2004). Ndc80C elimination could therefore affect CENP-E and RZZS recruitment indirectly by reducing kinetochore MPS1. Conversely, KNL1 recruits BUB1, also in an MPS1-dependent manner (Kiyomitsu *et al*, 2007; London *et al.*, 2012; Pagliuca *et al.*, 2009; Pereira *et al.*, 2018; Primorac *et al.*, 2013; Rodriguez-Rodriguez *et al.*, 2018; Shepperd *et al*, 2012; Silio *et al.*, 2015; Vleugel *et al.*, 2013; Yamagishi *et al.*, 2012). BUB1 further recruits BUBR1 (Overlack *et al.*, 2015), thus in turn contributing to CENP-E recruitment. Furthermore, BUB1 has been proposed to contribute more directly to RZZS recruitment (Caldas *et al.*, 2015; Zhang *et al*, 2015).

We reasoned that R^EE^ZZ is ideally suited to overcome these interdependencies, as it bypasses requirements for MPS1 activity and CENP-E. Thus, we adopted the experimental scheme in Figure 6A to assess if electroporated R^EE^ZZ was recruited to kinetochores in cells depleted of CENP-E and of either KNL1 or Ndc80C. Co-depletion of KNL1 and CENP-E resulted in a very severe depletion of mCherry-R^EE^ZZ from kinetochores (Figure 6B-C). Conversely, co-depletion of Ndc80C and CENP-E caused only a reduction of mCherry-R^EE^ZZ. As the extent of the reduction was comparable to that affecting KNL1 levels after Ndc80C depletion, these results seem to confirm a more prominent role of KNL1 in RZZ recruitment (Figure 6B-C and Figure 6-Supplement 1D-E). Further treatment with Reversine to inhibit MPS1 led to essentially complete disappearance of mCherry-R^EE^ZZ from kinetochores both in CENP-E/KNL1 and in CENP-E/Ndc80C co-depleted cells (Figure 6-Supplement 1G-H. Note that endogenous RZZ was also depleted from kinetochores in these conditions, see Figure 6-Supplement 1F and I).

Collectively, these results emphasize the role of Knl1C, or of a protein associated with Knl1C, as crucial kinetochore receptor of mCherry-R^EE^ZZ, while Ndc80C may play an only marginal role as a direct recruiter. The strong Reversine sensitivity of mCherry-R^EE^ZZ in cells depleted of CENP-E and Ndc80C suggests that even in absence of Ndc80C, considered the main MPS1 recruiter (see above), MPS1 activity continues to contribute to RZZ recruitment. As BUB1 is recruited to KNL1 in an MPS1-dependent manner (see Introduction), we assessed if depletion of KNL1 affected RZZS recruitment exclusively through BUB1. For this, we co-depleted CENP-E and BUB1 and compared the levels of residual mCherry-R^EE^ZZ (and of RZZ^WT^ as control; Figure 6D-G) at kinetochores to those observed when CENP-E and KNL1 were co-depleted instead (Figure 6-Supplement 1J-L). BUB1 and CENP-E co-depletion was significantly less effective than KNL1/CENP-E co-depletion in preventing RZZS kinetochore recruitment (Figure 6G-I). These results indicate that KNL1 may contribute to kinetochore recruitment of RZZS not only through BUB1, but possibly also by providing an additional binding site. Addition of Reversine to BUB1/CENP-E co-depleted cells further reduced the levels of mCherry-R^EE^ZZ (Figure 6F-I), suggesting that the effects of MPS1 inhibition on RZZS recruitment are not limited to BUB1. Furthermore, small residual levels of mCherry-R^EE^ZZ in KNL1/CENP-E co-depletion were almost entirely eliminated upon addition of MPS1 inhibitor (Figure 6F-G). Similar conclusions were reached upon co-depletion of BUB1 and Ndc80C, which caused a very strong reduction of endogenous RZZS and CENP-E. Also in this case, further addition of Reversine cleared kinetochores of these proteins (Figure 6-Supplement 2A-C). On the basis of these results, we conclude that RZZS likely also interacts weakly with the Ndc80C and that MPS1 facilitates interactions with both the Ndc80C and KNL1.

## Discussion

We discovered an interaction of CENP-E and RZZS that controls the recruitment and activation of DD at the kinetochore. By harnessing various separation of function mutants, we brought to light several new points of contact of the corona with the kinetochore. After our new characterization, the corona assembly plan of Figure 1B is in need of significant revision (Figure 7A). Our main results can be summarized in the following points.

**Figure 7.**
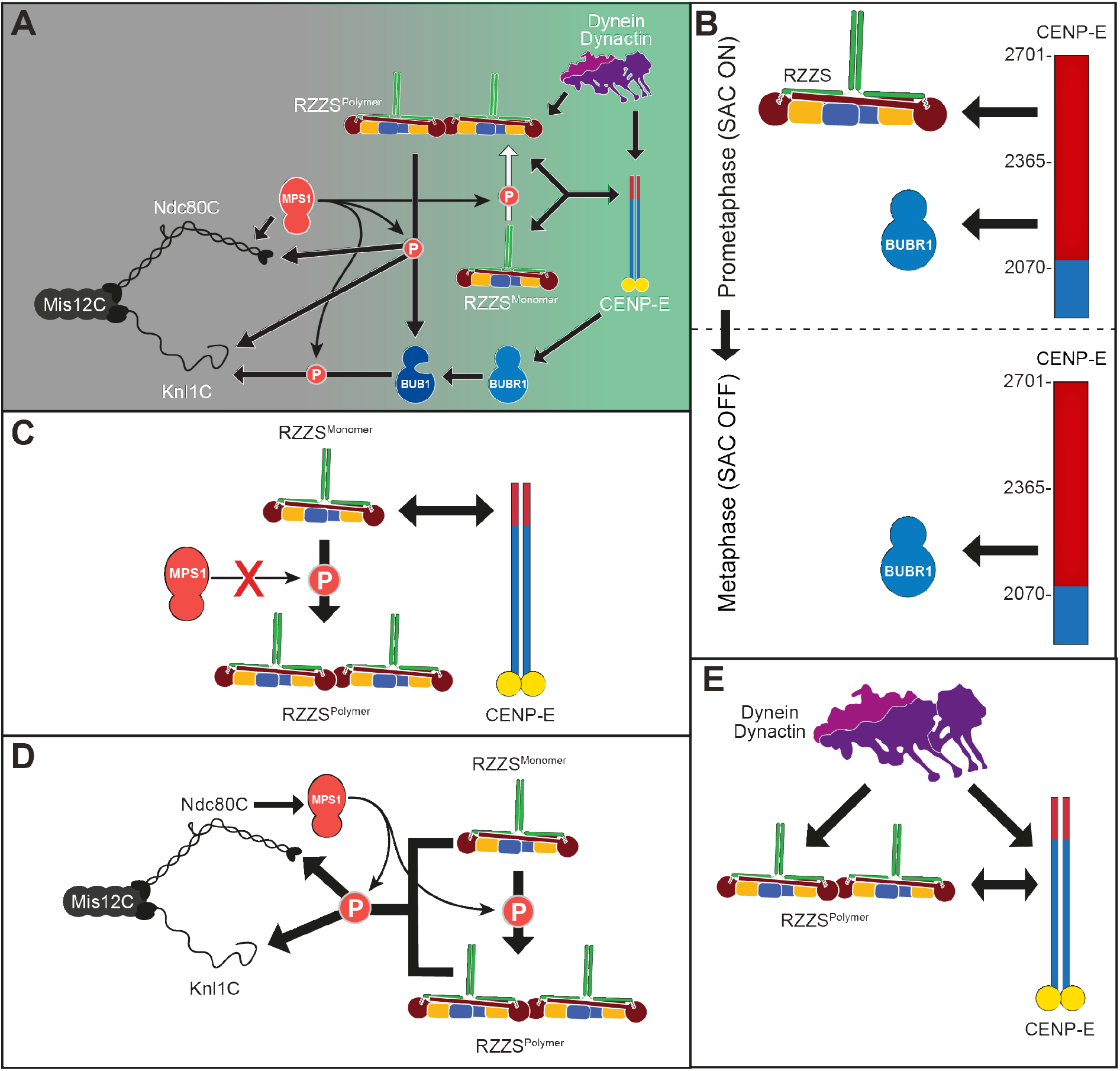
Summary of main results. (**A**) Revised drawing depicting the hierarchical organization of outer kinetochore and kinetochore corona components after our study. Compare with Figure 1B. Thick arrows indicate recruitment of a protein to the protein indicated by the arrowhead. Thin arrows indicate phosphorylation. The white arrow indicates polymerization. (**B**-**E**) Depiction of individual interactions in the order in which they are presented in the Discussion.

First, BUBR1 and RZZS play a highly prominent role in CENP-E recruitment. The interaction of CENP-E with BUBR1 had been described earlier (Ciossani *et al.*, 2018; Legal *et al.*, 2020; Mao *et al.*, 2003; Mao *et al.*, 2005). This interaction of CENP-E is clearly orthogonal to that with the RZZS complex, as it engages a different region of CENP-E and is sufficient for recruitment of CENP-E to kinetochores when the RZZS-binding site is mutated. The interaction with BUBR1 is dispensable for kinetochore recruitment of CENP-E in prometaphase, when CENP-E is clearly identified in the corona, but becomes essential for kinetochore recruitment of CENP-E after corona shedding, an event that coincides with biorientation (Figure 7B). At present, we are unable to conclude whether CENP-E is simultaneously bound to BUBR1 and the RZZS complex. The observation that BUBR1 does not integrate into the corona may suggest the existence of two distinct pools of CENP-E. However, CENP-E is highly elongated (Kim *et al.*, 2008), and its long axis may transverse the depth of the kinetochore, interacting with the RZZS complex within the corona but extending further inside the kinetochore to interact with BUBR1.

Constructs carrying BUBR1^Mut^ and RZZS^Mut^ were unable to localize to kinetochores. Conversely, depletion of BUBR1 and RZZS largely reduced kinetochore CENP-E, but did not completely eliminate it. The residual levels of CENP-E may reflect incomplete depletion of RZZ and BUBR1 or the existence of a third, currently unknown receptor. Below, we discuss our evidence that this unknown receptor may be the Ndc80C. In line with at least two earlier studies (Martin-Lluesma *et al.*, 2002; Sharp-Baker & Chen, 2001), we did not find evidence for an involvement of MAD1 in the recruitment of CENP-E (Akera *et al.*, 2015). Specifically, we found that a condition that largely reduced kinetochore MAD1 did not grossly affect CENP-E recruitment.

Second, RZZS and CENP-E are co-dependent for their own kinetochore recruitment (Figure 7C). In previous work, we and others had shown that depletion of CENP-E did not prevent corona assembly (Ciossani *et al.*, 2018; Martin-Lluesma *et al.*, 2002). Here, we confirmed this conclusion with a cell line that allows acute depletion of CENP-E (Owa & Dynlacht, 2021). However, depletion of CENP-E caused complete depletion of RZZS from kinetochores when combined with inhibition of MPS1 kinase. MPS1 has been previously shown to promote corona expansion by phosphorylating ROD (Raisch *et al.*, 2022; Rodriguez-Rodriguez *et al.*, 2018). These results indicate to us that MPS1, through corona expansion, through the facilitation of a direct interaction, or through both, promotes binding of RZZS to one or more kinetochore receptors, in addition to CENP-E. We refer to this receptor or receptors as “core kinetochore receptor” of RZZS, and comment on it below.

Third, MPS1, in addition to an established role in corona expansion through ROD phosphorylation (Raisch *et al.*, 2022; Rodriguez-Rodriguez *et al.*, 2018), indeed has a distinct function in promoting binding of RZZS to its core kinetochore receptor (Figure 7D). This additional function of MPS1 became evident when CENP-E depletion was combined with inhibition of corona expansion by means other than inhibiting MPS1, including expressing mutants of ROD or Zwilch, or depleting Spindly. Like MPS1 inhibition, these conditions did not prevent RZZS (or RZZ) kinetochore recruitment. Further inhibition of MPS1, however, abrogated RZZS or RZZ localization. Thus, lack of corona expansion with these alternative perturbations does not fully recapitulate the effects of inhibiting MPS1, indicating that MPS1, in addition to corona expansion, promotes binding of RZZS to its core kinetochore receptor. Importantly, the R^EE^ZZ mutant was insensitive to MPS1 inhibition and CENP-E depletion, and expanded a robust corona. Because R^EE^ZZ interacts with kinetochores when corona expansion and MPS1 are both inhibited (and in absence of CENP-E), we suspect that phosphorylation of Thr13 and Ser15, in addition to causing corona expansion, also mediates an interaction with the kinetochore, a question that will require more detailed investigations.

Fourth, BUB1, KNL1, and Ndc80C contribute to the establishment of the core kinetochore receptor of RZZS (also in Figure 7D). MPS1 promotes robust kinetochore recruitment of BUB1, which is in turn important for BUBR1 recruitment (Krenn *et al.*, 2014; London *et al.*, 2012; Meadows *et al.*, 2011; Overlack *et al.*, 2015; Primorac *et al.*, 2013; Vleugel *et al.*, 2013; Yamagishi *et al.*, 2012). It is therefore natural to turn to BUB1 as a plausible core kinetochore receptor of the RZZS, as previously done (Caldas *et al.*, 2015; Zhang *et al.*, 2015). CENP-E and BUB1 co-depletion, however, affected RZZS recruitment only marginally, and did not phenocopy the combination of CENP-E depletion and MPS1 inhibition. Thus, BUB1 depletion and MPS1 inhibition are not equivalent towards the outcome of recruiting RZZS in absence of CENP-E. Conversely, CENP-E and KNL1 co-depletion, but not CENP-E and Ndc80C co-depletion, counteracted RZZS recruitment to a decidedly more significant extent. When only KNL1 was depleted, substantial levels of residual RZZS were observed, in agreement with previous work (Caldas *et al.*, 2015). Because this residual pool of RZZS largely depended on CENP-E, recruitment of CENP-E, at least for the branch that does not depend on BUBR1, must be independent of KNL1. As co-depletion of KNL1 and Ndc80C prevented CENP-E and RZZS recruitment entirely, we reason that CENP-E, in addition to binding to BUBR1 and RZZS, likely also establishes an additional contact with Ndc80C that contributes to holding it at kinetochores when KNL1 has been depleted. We therefore speculate that the small residual levels of RZZS observed in cells co-depleted of KNL1 and CENP-E may also reflect an interaction with Ndc80C. Our data suggest also this interaction to require MPS1. Additional studies will be needed to identify the complete repertoire of binding determinants to allow us to build a complete model for the corona recruitment mechanism.

Fifth, the co-dependent localization of RZZS and CENP-E is especially meaningful for full DD recruitment to the kinetochore (Figure 7E). Previously, we have garnered evidence that Spindly, as a DD adaptor at the kinetochore, requires a kinetochore stimulus to “open up” from an auto-inhibited conformation (d’Amico *et al.*, 2022). However, Spindly mutants that we suspect to be in an open conformation based on their ability to interact with the pointed-end complex of Dynactin do not rescue the reduction of DD caused by CENP-E depletion. Thus, while we cannot conclude or exclude that CENP-E is the factor that “opens” Spindly at the kinetochore, we can argue that it is additionally required downstream of Spindly opening, possibly through a direct interaction stabilizing DD at the kinetochore. In the future, we will try to use biochemical reconstitution to gain further support for this model and identify the detailed determinants of this regulation. Our results also imply that depletion of CENP-E, by causing a significant co-depletion of DD, may have distinct and more pervasive effects on chromosome alignment than the sole inhibition of CENP-E with small molecule inhibitors (Ohashi *et al*, 2015; Wood *et al*, 2010).

While until now the corona has been primarily viewed as a platform for the coordination of Dynein motility and spindle assembly checkpoint activity, our results indicate that it should be instead regarded as an extended adaptor for opposite-polarity motors. The corona combines DD and CENP-E in a single integrated complex capable of bidirectional transport of chromosomes as cargo. Integration of minus- and plus-end-directed microtubule motor activity may be a common theme of transport for other categories of cargo, including lysosomes or secretory vesicles among others (Canty *et al*, 2021; Celestino *et al*, 2022; Fenton *et al*, 2021; Kendrick *et al*, 2019). TRAK1/2 adaptors, for instance, promote the incorporation of kinesin-1 (Kif5) and DD in a single complex (Canty *et al.*, 2021; Fenton *et al.*, 2021). Kinesin-1 binds a region of TRAK that contains the CC1 box, a DD binding region that we have also implicated in the intramolecular regulation of Spindly (d’Amico *et al.*, 2022; Randall *et al*, 2013). CENP-E may contribute to the activation of Spindly in a similar manner, in addition to a more direct role in DD binding. Collectively, our studies and the recent observations on the interactions of DD adaptors with kinesins seem to suggest that in addition to bringing Dynein and Dynactin into the same complex (Carter *et al.*, 2016; Olenick & Holzbaur, 2019; Reck-Peterson *et al.*, 2018), a more general function of adaptors and associated proteins is to bring DD and cognate kinesins into the same complex.

What triggers corona shedding upon end-on attachment is a question of great interest. Importantly, shedding will also deplete kinetochores of the MAD1:MAD2 core complex crucially required for checkpoint signaling, thus beginning checkpoint silencing (De Antoni *et al*, 2005; Fava *et al.*, 2011; Luo *et al*, 2018; Maldonado & Kapoor, 2011). Autoinhibition of kinesin-1 has been shown to facilitate the initiation of Dynein cargo transport (Qiu *et al*, 2023). Similarly, the transition to pole-directed transport of kinetochore proteins during shedding may reflect initiation of dynein cargo transport when the activity of CENP-E becomes suppressed. The relief from autoinhibition that prefigures activation of CENP-E motor activity may require phosphorylation by mitotic kinases, including MPS1 and CDK1 (Espeut *et al.*, 2008; Nousiainen *et al*, 2006). Aurora A and Aurora B, kinases residing primarily at spindle poles and centromeres, respectively, may further contribute to CENP-E activation by phosphorylating Thr422. This residue is encompassed within a binding motif for protein phosphatase 1 (PP1), whose phosphorylation counteracts PP1 recruitment (Egloff *et al*, 1997; Kim *et al.*, 2010; Liu *et al*, 2010). Upon dephosphorylation, docking of PP1 to T422 may promote further CENP-E dephosphorylation and its subsequent autoinhibition (Kim *et al.*, 2010). This, in turn, may initiate shedding. This model is attractive because Aurora B kinase has an established role in bi-orientation and is believed to be regulated by forces exerted by microtubules as they bind the kinetochore (Krenn & Musacchio, 2015; Lampson & Grishchuk, 2017). We note that the corona appeared to disassemble normally in presence of the phosphomimetic R^EE^ ZZ mutant, suggesting that preventing dephosphorylation of the MPS1 sites may not be sufficient for retaining the corona. Changes in MPS1 or Aurora B activity upon bi-orientation, however, are likely to be relevant for shedding. Dephosphorylation of MPS1 sites may cause a reduction in the binding affinity of RZZS for its core kinetochore receptor KNL1, which we show to depend on MPS1 activity. Dephosphorylation of T422 of CENP-E and other Aurora B sites, by causing conformational changes in CENP-E, may also affect its ability to hold onto the corona.

The identification of binding determinants is prerequisite for the dissection of their dynamic regulation and for the generation of adequate separation-of-function mutants. In this study, we have identified several new interactions that are essential for the establishment of the kinetochore corona. Understanding the regulation of these interactions will contribute to dissect the basis for the coordination of chromosome biorientation and checkpoint silencing. Our work sets the stage for future investigations of the dauntingly complex processes that enforce correct chromosome segregation.

## Methods

### Materials and Methods

#### Mutagenesis and cloning

The codon-optimized cDNA of CENP-E (Q02224) was synthesized at GeneWiz and subcloned in pLIB-EGFP and pET-EGFP, modified versions of, respectively, the pLIB (Weissmann *et al*, 2016) and pET-28 vector for expression of proteins with N-terminal PreScission-cleavable His_6_-EGFP-tag. Mutations were introduced by site-directed mutagenesis and Gibson assembly (Gibson *et al*, 2009) and verified by Sanger sequencing (Microsynth Seqlab). The Spindly constructs were generated as previously described (d’Amico *et al.*, 2022) and subcloned in pLIB with an N-terminal His_6_-mCherry-tag.

#### Expression and purification of RZZ, GST-Spindly and mCherry-Spindly constructs

The RZZ complex was expressed and purified using the biGBac system (Weissmann *et al.*, 2016), with a mCherry-tag on the N-terminus of the ROD subunit, as previously described (Raisch *et al.*, 2022; Sacristan *et al.*, 2018). All Spindly constructs, except for Spindly^33-605^, were expressed using the biGBac system. Spindly^33-605^ was expressed in *Escherichia coli (E. coli)* and purified as described (d’Amico *et al.*, 2022). The Baculovirus was generated in Sf9 cells to infect Tnao38 cells, which were grown for 72 hours at 27° C before harvesting. The pellet was washed with PBS, snap-frozen and stored at -80° C. Purification of the Spindly constructs was performed as previously described (d’Amico *et al.*, 2022) The pellet was resuspended in lysis buffer (50 mM HEPES pH 8.0, 250 mM NaCl, 30 mM imidazole, 2 mM Tris(2-carboxyethyl)phosphine (TCEP)) supplemented with protease inhibitor, 1 mM PMSF, DNaseI and lysed by sonication. The lysate was clarified by centrifugation for 45 minutes at 88,000 g, sterile filtered and loaded onto a HisTrap HP column (Cytiva). Subsequently, the column was washed with at least 20 column volumes lysis buffer. Elution was performed with lysis buffer supplemented with 300 mM imidazole. The eluate was diluted 1:5 in no salt buffer (50 mM HEPES pH 8.0, 2 mM TCEP), and applied to a 6 ml Resource Q anion exchange column (Cytiva). The protein was eluted with a 50-500 mM NaCl gradient, and fractions were analyzed by SDS-PAGE. Fractions containing the protein of interest were pooled and concentrated. Finally, the protein was loaded onto a Superdex 200 16/60 pre-equilibrated in SEC buffer (50 mM HEPES pH 8.0, 250 mM NaCl, 2 mM TCEP). Peak fractions containing the protein of interest were analyzed by SDS-PAGE, concentrated, snap-frozen, and stored at -80° C until further usage.

#### Expression and purification of EGFP-CENP-E constructs

Expression of CENP-E^2070C^ wild-type and mutants and CENP-E^2070-2365^ was carried out in insect cells using the biGBac system. The Baculoviruses were generated in Sf9 cells to infect Tnao38 cells for 72 hours at 27 °C before harvesting. The pellet was washed with PBS, snap-frozen and stored at -80° C. For purification, CENP-E was resuspended in lysis buffer (50 mM HEPES pH 8.0, 250 mM NaCl, 5% (w/v) glycerol, and 1 mM TCEP) supplemented with protease inhibitor, 1 mM PMSF, DNaseI and lysed by sonication. The lysate was clarified by centrifugation for 45 minutes at 88,000 g, sterile filtered and loaded onto a HisTrap HP column (Cytiva). Elution was performed with lysis buffer supplemented with 300 mM imidazole. Subsequently, the CENP-E^2070C^ mutants were diluted 1:5 in low salt buffer (50 mM HEPES pH 8.0, 50 mM NaCl, 5% (w/v) glycerol, 1 mM TCEP), and applied to a 6 ml Resource Q anion exchange column (Cytiva). The protein was eluted with a 50-500 mM NaCl gradient, and peak fractions were analyzed by SDS-PAGE. Fractions containing the protein of interest were pooled and concentrated. The concentrated sample was loaded onto a Superdex 200 16/60 pre-equilibrated in SEC buffer (50 mM HEPES pH 8.0, 250 mM NaCl, 5% (w/v) glycerol, 1 mM TCEP). Finally, peak fractions containing the protein of interest were analyzed by SDS-PAGE, concentrated, snap-frozen and stored at -80° C until further usage. CENP-E^2070C^ wild-type was loaded on a Superdex 200 16/60 directly after affinity purification. Peak fractions containing the protein of interest were analyzed by SDS-PAGE, concentrated, snap-frozen, and stored at -80° C until further usage.

EGFP-CENP-E^2366C^ (WT and mutants) were expressed in *E. coli* BL21 (DE3) RP plus cells grown in Terrific-Broth (TB) at 37°C to A_600_ = 2 and then induced for 16 h at 17°C with 0.25 mM isopropyl-beta-D-thiogalactopyranoside (IPTG). Cells were collected by centrifugation, washed in PBS, and then frozen at -80°C. Cell pellets were resuspended in lysis buffer (50 mM sodium phosphate pH 7.0, 5% (w/v) glycerol, 2 mM β-mercaptoethanol and 500 mM NaCl) supplemented with protease inhibitor, lysed by sonication, and cleared by centrifugation at 70,000 g at 4°C. The supernatant was filtered and loaded on a 5 ml HisTrap FF column (GE Healthcare) equilibrated in lysis buffer. After washing with lysis buffer and 75 mM imidazole, the proteins were eluted with 500 mM imidazole. Proteins were concentrated with a 30 kDa cutoff Amicon concentrator (Millipore), and gel-filtered on a Superose 6 10/30 (GE Healthcare) equilibrated in SEC buffer (50 mM HEPES pH 7.0, 200 mM NaCl, 5% (w/v) glycerol, 1 mM TCEP).

Expression and purification of the kinase domain of BUBR1 (BUBR1^KD^) construct was carried out as previously reported (Breit *et al.*, 2015).

#### Analytical Size Exclusion Chromatography (SEC)

Binding assays of Spindly with CENP-E^2070C^and PE complex were performed under isocratic conditions on a Superdex 200 15/50 pre-equilibrated in SEC buffer (50 mM HEPES pH 8.0, 150 mM NaCl, 2 mM TCEP) at 4° C on an ÄKTAmicro system. Elution of proteins was monitored at 280 nm. 50 µl fractions were collected and analyzed by SDS-PAGE. To assess complex formation proteins were mixed at the indicated concentrations in 60 µl SEC buffer and incubated for at least 1 hour on ice before the SEC assay was performed.

Binding assays of CENP-E and BUBR1 were carried out by mixing 16 µM CENP-E proteins with 8 µM BUBR1^KD^ in a final volume of 30 µl. Analytical size-exclusion chromatography was carried out under isocratic conditions on a Superose 6 5/150 or Superdex 200 5/150 (GE Healthcare) equilibrated with SEC buffer (50 mM HEPES pH 8.0, 100 mM NaCl, 5% (w/v) glycerol and 0.5 mM TCEP) at 4° C on an ÄKTAmicro system. Protein elution was monitored at 280 nm and 50 µl fractions were collected and analyzed by SDS-PAGE followed by Coomassie Blue staining.

#### RZZ-Spindly filament formation and imaging

RZZS filaments were formed essentially as described in (d’Amico *et al.*, 2022). 4 µM mCherry-RZZ and 8 µM prefarnesylated Spindly was incubated overnight at room temperature in the presence of 1 µM GST-MPS1 in M-buffer (50 mM HEPES pH 7.5, 100 mM NaCl, 1 mM MgCl_2_, 1 mM TCEP). Flow chambers were assembled by applying two strips of double-sided tape on a glass slide and then placing a standard coverslip on top. The sample was diluted 1:8 and EGFP-CENP-E constructs were added to a final concentration of 1 µM, promptly flowed into a flow chamber and imaged. Imaging was performed on a 3i Marianas system at 100x magnification. Sample images were acquired as five-stacks of z-sections at 0.27 µm, maximum-projected on the z-axis, and processed in Fiji.

#### RZZ-Spindly rings formation and imaging

RZZS rings were formed by incubating 4 µM mCherry-RZZ with 8 µM prefarnesylated mCherry-Spindly in the presence of 1 µM GST-MPS1 in M-buffer. Flow chambers were assembled by placing two strips of double-sided tape on a glass slide and a pre-cleaned high-performance cleanroom cleaned 1.5H coverslip (Nexterion) on top. The flow chamber was maintained at room temperature for all subsequent steps. The chamber was first equilibrated in S-buffer (50 mM HEPES pH 8, 200 mM NaCl, 2 mM TCEP). Anti-mCherry GST-tagged nanobodies (Addgene #70696, (Katoh *et al*, 2016)) were expressed and purified according to published protocol (Katoh *et al.*, 2016). The nanobody was diluted in 20 µl S-buffer to a final concentration of 1 µM, flowed through the chamber, and incubated for 5-10 minutes. The chamber was then washed with 20 µl S-buffer with 1% pluronic F-127 and incubated for 5 minutes. Following passivation, a wash was performed with 50 µl S-buffer with 0.1 mg/ml BSA (A-buffer) to saturate aspecific binding sites. The rings were diluted 1:100 in 20 µl A-buffer, flowed into the chamber, incubated for 5 minutes, and washed with A-buffer. EGFP-CENP-E and BUBR1 constructs were mixed and diluted to the specified concentrations in 20 µl A-buffer and flowed into the chamber, which was then promptly sealed and moved to the microscope for imaging. Imaging was performed on a VisiTIRF microscope in the 488 nm and 561 nm laser channels, at 100x magnification, in TIRF mode. Single images were taken with exposures of 100 or 200 ms. Images were visualized and cropped using Fiji.

##### Molecular modeling

CENP-E and Spindly predictions were generated using AlphaFold Multimer (Evans *et al*, 2021; Jumper *et al.*, 2021) as previously described (d’Amico *et al.*, 2022).

##### Mammalian plasmids

All mammalian plasmids were derived from pCDNA5/FRT/TO-EGFP-IRES, a previously modified version (Krenn *et al*, 2012) of the pCDNA5/FRT/TO vector (Invitrogen). To generate N-terminally-tagged EGFP-CENP-E constructs, the CENP-E sequence was obtained by PCR and subcloned in-frame with the EGFP-tag. All CENP-E constructs are RNAi-resistant and validated by Sanger sequencing (Microsynth Seqlab).

##### Cell lines

Parental Flp-In T-REx DLD-1 osTIR1 cells were a kind gift from D. C. Cleveland (University of California, San Diego, USA). The hTERT-immortalized retinal pigment epithelial (hTERT-RPE-1) cell line in which both CENP-E alleles are C-terminally tagged with an AID and a 3xFLAG tag has been described (Owa & Dynlacht, 2021). The cell line expresses the plant E3 ubiquitin ligase osTIR1, which was stably integrated into the genome. Degradation of the endogenous CENP-E is achieved through addition of 500 μM IAA (Sigma-Aldrich) (Owa & Dynlacht, 2021).

##### Cell culture

HeLa, DLD-1 and RPE-1 cells were grown in Dulbecco’s Modified Eagle’s Medium (DMEM; PAN Biotech) supplemented with 10 % tetracycline-free FBS (PAN Biotech), and L-Glutamine (PAN Biotech). Cells were grown at 37°C in the presence of 5 % CO_2_.

##### Generation of stable cell lines

Stable Flp-In T-REx DLD-1 osTIR1 cell lines were generated using FRT/Flp recombination. CENP-E constructs were cloned into a pCDNA5/FRT/TO-EGFP-IRES plasmid and co-transfected with pOG44 (Invitrogen), encoding the Flp recombinase, into cells using X-tremeGENE (Roche) according to the manufacturer’s instructions. Subsequently, cells were selected for 2 weeks in DMEM supplemented with hygromycin B (250 μg/ml; Thermo Fisher Scientific) and blasticidin (4 μg/ml; Thermo Fisher Scientific). Single-cell colonies were isolated, expanded and expression of the transgenes was checked by immunofluorescence microscopy and immunoblotting analysis. Gene expression was induced by addition of 0.01–0.3 µg/ml doxycycline (Sigma-Aldrich).

##### RNAi and drug treatment

Depletion of endogenous proteins was achieved through transfection of single small interfering RNA (siRNA) with RNAiMAX (Invitrogen) according to manufacturer’s instructions. Following siRNAs treatments were performed in this study: 50 nM siBUB1 (Dharmacon, 5’-GGUUGCCAACACAAGUUCU3’) for 24 h, 100 nM siBUBR1 (Dharmacon, 5’-CGGGCAUUUGAAUAUGAAA-3’) for 24 h, 60 nM siCENP-E (Dharmacon, 5’-AAGGCUACAAUGGUACUAUAU-3’) for 24 h, 50 nM siCENP-F (Dharmacon, 5’-CAAAGACCGGUGUUACCAAG-3’; 5’-AAGAGAAGACCCCAAGUCAUC-3’) for 24 h, siKNL1 (Invitrogen, HSS183683 5’-CACCCAGUGUCAUACAGCCAAUAUU-3’; HSS125942 5’-UCUACUGUGGUGGAGUUCUUGAUAA-3’; CCCUCUGGAGGAAUGGUCUAAUAAU-3’) for 24 h, siSpindly (Sigma-Aldrich, 5′-GAAAGGGUCUCAAACUGAA-3′) for 48 h, siNdc80C (Sigma-Aldrich, siHec1 5’-GAGUAGAACUAGAAUGUGA-3’; siSpc24 5’-GGACACGACAGUCACAAUC-3’; siSpc25 5’-CUACAAGGAUUCCAUCAAA-3’) for 48 h, 100 nM siZwilch (SMART pool from Dharmacon, L-019377 -00-0005) for 72 h.

Unless indicated, Nocodazole (Sigma-Aldrich) was used at 3.3 µM, RO3306 (Calbiochem) at 9 µM, MG-132 (Calbiochem) at 10 µM, and Reversine (Cayman Chem.) at 500 nM.

##### Electroporation of recombinant protein into human cells

Recombinant mCherry-RZZ, mCherry-Spindly, or EGFP-CENP-E constructs were electroporated as previously described (Alex *et al.*, 2019) using the Neon Transfection System Kit (Thermo Fisher Scientific). HeLa cells were trypsinized, washed with PBS, and resuspended in electroporation buffer R (Thermo Fisher Scientific). Recombinant protein was added to a final concentration of 7 μM in the electroporation slurry and electroporated by applying two consecutive 35-ms pulses with an amplitude of 1000 V. Control cells were electroporated with mCherry or electroporation buffer, respectively. The electroporated sample was subsequently added to 15 ml of prewarmed PBS, centrifuged at 1500 rpm for 5 minutes, and trypsinized for 5 minutes to remove noninternalized extracellular protein. After two additional PBS washing and centrifugation steps, the cell pellet was resuspended in prewarmed DMEM and seeded in a 6-well plate. Following an 8-hour recovery, cells were treated with 9 µM RO3306 (Calbiochem) for 15 hours. Subsequently, cells were released into mitosis in presence of 3.3 µM Nocodazole for 1 hour before fixation for immunofluorescence or harvesting for immunoblotting.

##### Immunofluorescence

Cells were grown on coverslips pre-coated with poly-L-lysine (Sigma-Aldrich). Before fixation, cells were pre-permeabilized with 0.5% Triton X-100 solution in PHEM (Pipes, HEPES, EGTA, MgCl_2_) buffer supplemented with 100 nM microcystin for 5 minutes before fixation with 4% PFA in PHEM for 20 minutes. For Dynactin-p150^glued^ staining the cells were initially fixated with 4% PFA in PHEM for 5 minutes and then permeabilized for 10 minutes with PHEM supplemented with 0.5% Triton-X-100. After blocking with 5% boiled goat serum (BGS) in PHEM buffer for 1 hour, cells were incubated for 2 hours at room temperature with the following primary antibodies: BUB1 (mouse, Abcam, ab54893, 1:400), BUBR1 (rabbit, Thermo Scientific #720297, 1:1000), CENP-C (guinea pig, MBL, #PD030, 1:1000), CENP-E (mouse, Abcam, ab5093, 1:200), CENP-F (rabbit, Novus NB500-101, 1:300), CREST/anti-centromere antibodies (human, Antibodies, Inc., 1:200), Dynactin-p150^glued^ (mouse, BD Trans. Lab., #610473, 1:150), Hec1 (mouse, Clone 9G3.23, Gene-Tex, Inc., 1:1000), KNL1 (rabbit, made in-house, #SI0787, 1:750), MAD1 labeled with DyLight488 (mouse, made in-house, Clone BB3-8, 1:200), Spindly (rabbit, Bethyl, #A301-354A, 1:1000), Zwilch (rabbit, made in-house, #SI520, 1:900) diluted in 2.5% BGS-PHEM supplemented with 0.1% Triton-X-100. Subsequently, cells were incubated for 1 hour at room temperature with the following secondary antibodies (all 1:200 in 2.5% BGS-PHEM supplemented with 0.1% Triton-X-100): Goat anti-mouse Alexa Fluor 488 (Invitrogen A A11001), goat anti-mouse Rhodamine Red (Jackson Immuno Research 115-295-003), donkey anti-rabbit Alexa Fluor 488 (Invitrogen A21206), donkey anti-rabbit Rhodamine Red (Jackson Immuno Research 711-295-152), goat anti-human Alexa Fluor 647 (Jackson Immuno Research 109-603-003), goat anti-guinea pig Alexa Fluor 647 (Invitrogen A-21450). All washing steps were performed with PHEM supplemented with 0.1% Triton-X-100 (PHEM-T) buffer. DNA was stained with 0.5 μg/ml DAPI (Serva) and Mowiol (Calbiochem) was used as mounting media.

##### Cell imaging

Cells were imaged at room temperature using a spinning disk confocal device on the 3i Marianas system equipped with an Axio Observer Z1 microscope (Zeiss), a CSU-X1 confocal scanner unit (Yokogawa Electric Corporation, Tokyo, Japan), 100 × /1.4NA Oil Objectives (Zeiss), and Orca Flash 4.0 sCMOS Camera (Hamamatsu). Alternatively, cells were imaged using a 60x oil immersion objective lens on a DeltaVision deconvolution microscope (GE Healthcare, UK) equipped with an IX71 inverted microscope (Olympus, Japan), a PLAPON ×60/1.42 numerical aperture objective (Olympus) and a pco.edge sCMOS camera (PCO-TECH Inc., USA). Confocal images were acquired as z sections at 0.27 μm (using Slidebook Software 6 from Intelligent Imaging Innovations). Images were converted into maximal intensity projections, converted into 16-bit TIFF files, and exported. Automatic quantification of single kinetochore signals was performed using the software Fiji with background subtraction. Measurements were exported in Excel (Microsoft) and graphed with GraphPad Prism 9.0 (GraphPad Software). Statistical analysis was performed with a nonparametric t-test comparing two unpaired groups (Mann-Whitney test). Symbols indicate: n.s. = p > 0.05, ∗ = p ≤ 0.05, ∗∗ = p ≤ 0.01, ∗∗∗ = p ≤ 0.001, ∗∗∗∗ = p ≤ 0.0001. Figures were arranged using Adobe Illustrator 2022.

##### Immunoblotting

Mitotic cells were collected via shake-off and resuspended in lysis buffer (150 mM KCl, 75 mM HEPES pH 7.5, 1.5 mM EGTA, 1.5 mM MgCl_2_, 10% (w/v) glycerol, and 0.075 % NP-40 supplemented with protease inhibitor cocktail (Serva) and PhosSTOP phosphatase inhibitors (Roche)). After lysis, whole-cell lysates were centrifuged at 15,000 rpm for 30 minutes at 4°C. Subsequently, the supernatant was collected and resuspended in sample buffer for analysis by SDS-PAGE and Western blotting. The following primary antibodies were used: FLAG-tag (rabbit, Thermo Scientific, #PA1-984B, 1:1000), GFP (rabbit, made in-house, 1:3000), mCherry (mouse, Novus Biologicals, #NBP1-96752, 1:2000), α-tubulin (mouse monoclonal, Sigma-Aldrich; 1:8000). As secondary antibodies, anti-mouse or anti-rabbit (1:10000; Amersham, NXA931 and NA934) conjugated to horseradish peroxidase were used. After incubation with ECL Western blotting reagent (GE Healthcare), images were acquired with the ChemiDoc MP System (Bio-Rad) using Image Lab 6.0.1 software.

## Author contributions

**Conceptualization:** V.C., G.C., A.M.; **Investigation:** V.C., E.dA., G.C.; **Funding acquisition:** A.M.; **Project Administration:** A.M.; **Resources:** S.W., M.O., B.D.; **Supervision:** A.M. **Validation:** A.M., V.C.; **Visualization:** V.C., E.dA., G.C.; **Writing – original draft:** A.M., V.C.; **Writing – review & editing:** All authors

## Supplemental Information

Figure 1 – Supplement 1

Figure 2 – Supplements 1

Figure 2 – Supplements 2

Figure 3 – Supplement 1

Figure 4 – Supplement 1

Figure 5 – Supplements 1

Figure 5 – Supplements 2

Figure 6 – Supplements 1

Figure 5 – Supplements 2

## Acknowledgements

We are grateful to Ingrid Vetter for generating AF2 models of CENP-E and Spindly. We are also grateful to Stefano Maffini and Nico Schmidt for help with microscopy experiments and data analysis. We are also grateful to Sabina Colombo, Felix Ruhnow, and Thomas Surrey for helpful discussion and collaborative experiments not included in the manuscript. We also thank Jingchao Wu, Geert Kops, Susana Eibes, Marin Barisic, and Julie Welburn for communicating results prior to publication. V.C. and A.M. acknowledge funding from the DFG’s Collaborative Research Centre “Molecular Mechanisms of Cell State Transitions” (CRC 1430). A.M. also acknowledges the Max Planck Society, the European Research Council (ERC) Synergy Grant 951430 (BIOMECANET), the Marie-Curie Training Network DivIDE (project number 675737), and the CANTAR network under the Netzwerke-NRW program. B.D.D. was supported by grant 5R01GM120776-08 (NIH).

