## Supplemental Figures for "A mechanism that integrates microtubule motors of opposite polarity at the kinetochore corona"

### Supplemental Figure Legends

#### *Figure 1 – Supplement 1*

(**A-B**) Quantification of BUBR1 and Zwilch levels of cells treated as shown in Figure 1C-D. n refers to individually measured kinetochores. (**C**) Representative images showing the localization of $^{EGFP}$CENP-E$^{2070C}$ in prometaphase and metaphase cells. 32 h after cells were seeded, protein expression was induced through addition of 300 ng/mL doxycycline for 16 h before fixation. CENP-C was used to visualize kinetochores and DAPI to stain DNA. Scale bar: 5 µm. (**D**) Mitotic index analysis of stable DLD-1 cell lines expressing $^{EGFP}$CENP-E$^{2070C}$. 32 h after seeding, cells were treated with 300 ng/mL doxycycline or DMSO for 16 h before fixation. n refers to the number of analyzed cells. (**E**) Representative images showing the localization of $^{EGFP}$CENP-E$^{2070\text{-}2365}$ in prometaphase in presence or absence of BUBR1. BUBR1 RNAi treatment was performed with 100 nM siRNA and CENP-E RNAi treatment with 60 nM siRNA. 13 h after CENP-E and BUBR1 RNAi treatment cells were electroporated with recombinant $^{EGFP}$CENP-E$^{2070\text{-}2365}$. Following an 8 h recovery, cells were synchronized in G2 phase with 9 µM RO3306 for 15 h and then released into mitosis. Subsequently, cells were immediately treated with 3.3 µM Nocodazole for an additional hour. CENP-C was used to visualize kinetochores and DAPI to stain DNA. Scale bar: 5 µm. (**F-G**) Quantification of BUBR1 and EGFP levels at kinetochores of the experiment shown in panel E. n refers to individually measured kinetochores. (**H-I**) Analytical SEC binding assays between the BUBR1 kinase domain (KD) and different $^{EGFP}$CENP-E constructs. The profile of the complex is represented as a continuous line, and the individual CENP-E constructs with a dashed line. BUBR1: 8 µM, CENP-E constructs: 16 µM.

#### *Figure 2 – Supplement 1*

(**A**-**B**) Multiple sequence alignment showing the kinetochore binding region of CENP-E was generated in Jalview with the MAFFT algorithm. Residues are depicted according to CLUSTAL color code. Amino acid substitutions comprised in the BUBR1$^{Mut}$ and RZZS$^{Mut}$ are labeled in red above the sequence alignment. (**C**) Analytical SEC binding assays between the BUBR1 kinase domain (KD) and different $^{EGFP}$CENP-E$^{2070C}$ constructs. The complex run is represented as a continuous line, and the individual CENP-E constructs with a dashed line. BUBR1: 8 µM, CENP-E constructs: 16 µM. (**D**) AF2 Multimer model of CENP-E$^{2070C}$. Insets show the BUBR1$^{Mut}$ and RZZS$^{Mut}$ (a previously published BUBR1$^{Mut}$) (Legal *et al.*, 2020) and surrounding sequence. The main chain is depicted in blue, and mutated residues in red. (**E**) RZZS filament-binding assay showing recruitment of $^{EGFP}$CENP-E$^{2070C}$ constructs to $^{mCh}$RZZS filaments. Scale bar: 5 µm. (**F**) RZZS ring-binding assay showing recruitment of $^{EGFP}$CENP-E$^{2070C}$ constructs to $^{mCH}$RZZ$^{mCh}$S rings. Scale bar: 5 µm.

***Figure 2 – Supplement 2***

(**A**) Immunoblot of mitotic DLD-1 cells stably expressing different $^{EGFP}$CENP-E$^{2070C}$ constructs, treated as shown in (Figure 2A) and probed with the indicated antibodies. 50 µg of cleared lysate was used for each condition, and Tubulin is shown as a loading control. (**B**) Representative images showing the localization of different $^{EGFP}$CENP-E$^{2070C}$ constructs in prometaphase after depletion of CENP-E with 60 nM siRNA. 13 h after RNAi treatment cells were electroporated with recombinant $^{EGFP}$CENP-E$^{2070C}$ constructs as indicated. Following an 8 h recovery, cells were synchronized in G2 phase with 9 µM RO3306 for 15 h and then released into mitosis. Subsequently, cells were immediately treated with 3.3 µM Nocodazole for an additional hour. CENP-C was used to visualize kinetochores and DAPI to stain DNA. Scale bar: 5 µm. (**C**) Quantification of EGFP levels at kinetochores of the experiment shown in panel B. n refers to individually measured kinetochores. (**D**) Representative images showing the localization of different $^{EGFP}$CENP-E$^{2070\text{-}2365}$ constructs in prometaphase after depletion of CENP-E with 60 nM siRNA. 13 h after RNAi treatment cells were electroporated with recombinant $^{EGFP}$CENP-E$^{2070\text{-}2365}$ constructs as indicated. Following an 8 h recovery, cells were synchronized in G2 phase with 9 µM RO3306 for 15 h and then released into mitosis. Subsequently, cells were immediately treated with 3.3 µM Nocodazole for an additional hour. CENP-C was used to visualize kinetochores and DAPI to stain DNA. Scale bar: 5 µm. (**E**) Quantification of EGFP levels at kinetochores of the experiment shown in panel D. n refers to individually measured kinetochores. (**F**) Representative images showing the localization of different $^{EGFP}$CENP-E$^{2366C}$ constructs in prometaphase after depletion of CENP-E with 60 nM siRNA. 13 h after RNAi treatment cells were electroporated with recombinant $^{EGFP}$CENP-E$^{2366C}$ constructs as indicated. Following an 8 h recovery, cells were synchronized in G2 phase with 9 µM RO3306 for 15 h and then released into mitosis. Subsequently, cells were immediately treated with 3.3 µM Nocodazole for an additional hour. CENP-C was used to visualize kinetochores and DAPI to stain DNA. Scale bar: 5 µm. (**G**) Quantification of EGFP levels at kinetochores of the experiment shown in panel F. n refers to individually measured kinetochores. (**H-I**) Comparison of BUBR1 and Zwilch levels at kinetochores in Nocodazole and MG132 arrested cells. n refers to individually measured kinetochores.

***Figure 3 – Supplement 1***

(**A**) Schematic of the cell synchronization and imaging experiment shown in panel B. (**B**) Representative images showing the localization of Zwilch in prometaphase after depletion of CENP-E with 60 nM siRNA. 8 h after RNAi treatment, cells were synchronized in G2 phase with 9 µM RO3306 for 15 h and then released into mitosis. Subsequently, cells were immediately treated with 3.3 µM Nocodazole, 10 µM MG132 and, where indicated, with 500 nM Reversine, for an additional hour. CENP-C was used to visualize kinetochores and DAPI to stain DNA. Scale bar: 5 µm. (**C-D**) Quantification of Zwilch and CENP-E levels at kinetochores of the experiment shown in panel B. n refers to individually measured kinetochores. (**E**) Representative images showing the localization of Zwilch in prometaphase after depletion of CENP-E with 60 nM siRNA. 13 h after RNAi treatment cells were electroporated with electroporation buffer or $^{EGFP}$CENP-E$^{2366C}$. Following an 8 h recovery, cells were synchronized in G2 phase with 9 µM RO3306 for 15 h and then released into mitosis. Subsequently, cells were immediately treated with 3.3 µM Nocodazole, 10 µM MG132 and, where indicated, with 500 nM Reversine, for an additional hour. CENP-C was used to visualize kinetochores and DAPI to stain DNA. Scale bar: 5 µm. (**F-G**) Quantification of EGFP and Zwilch levels at kinetochores of the experiment shown in panel E. n refers to individually measured kinetochores. (**H**) Representative images showing the localization of MAD1 in cells treated as shown in Figure 3A. (**I**) Quantification of MAD1 levels at kinetochores of the experiment shown in panel H. n refers to individually measured kinetochores. (**J**) Representative images showing the localization of MAD1 after inhibition of MPS1. 32 h after seeding, cells were synchronized in G2 phase with 9 µM RO3306 for 15 h and then released into mitosis. Subsequently, cells were immediately treated with 3.3 µM Nocodazole, 10 µM MG132 and, where indicated, with 500 nM Reversine, for an additional hour. CENP-C was used to visualize kinetochores and DAPI to stain DNA. Scale bar: 5 µm. (**K-L**) Quantification of CENP-E and MAD1 levels at kinetochores of the experiment shown in panel J. n refers to individually measured kinetochores.

***Figure 4 – Supplement 1***

(**A**) AF2 Multimer model of Spindly$^{1-309}$ (d'Amico *et al.*, 2022) and multiple sequence alignment showing the CC2 region of Spindly was generated in Jalview. Residues are depicted according to CLUSTAL color code. Amino acid substitutions mutated in the Spindly$^{4A}$ construct are labeled in magenta above the sequence alignment. The inset in the AF2 model shows amino acids 275-306 of Spindly and surrounding sequence. The main chain is depicted in green and mutated residues in magenta. (**B-C**) Analytical SEC binding assays between the Dynactin-PE (brown) and $^{mCh}$Spindly constructs. The complex run is represented as a continuous line, and the individual Spindly constructs with a dashed line. PE: 3 µM, Spindly constructs: 16 µM. The control gels with Dynactin-PE alone are shared between panels B and C. (**D-E**) Analytical SEC binding assays between the CENP-E$^{2070C}$ and $^{mCh}$Spindly constructs. The complex run is represented as continuous line and the individual constructs with a dashed line. CENP-E: 20 µM, Spindly constructs: 16 µM. The control gels with $^{mCh}$Spindly$^{4A}$ alone are shared between panels C and E. (**F-G**) Quantification of Dynactin-p150$^{glued}$ and mCherry levels at kinetochores after depletion of CENP-E with 60 nM siRNA and Spindly with 50 nM siRNA. 13 h after CENP-E RNAi treatment cells were electroporated with electroporation buffer or recombinant $^{mCh}$Spindly constructs as indicated. Following an 8 h recovery, cells were synchronized in G2 phase with 9 µM RO3306 for 15 h and then released into mitosis. Subsequently, cells were immediately treated with 3.3 µM Nocodazole for an additional hour. n refers to individually measured kinetochores. (**H**) Representative images showing the localization of Dynactin monitored through the p150$^{glued}$ subunit after depletion of CENP-E with 60 nM siRNA and CENP-F with 60 nM. 8 h after RNAi treatment, cells were synchronized in G2 phase with 9 µM RO3306 for 15 h and then released into mitosis. Subsequently, cells were immediately treated with 3.3 µM Nocodazole for an additional hour. CENP-C was used to visualize kinetochores and DAPI to stain DNA. Scale bar: 5 µm. (**I-K**) Quantification of CENP-E, CENP-F and Dynactin-p150$^{glued}$ levels at kinetochores of the experiment shown in panel H. n refers to individually measured kinetochores.

***Figure 5 – Supplement 1***

(**A**) Representative images showing cells electroporated with the indicated $^{mCh}$RZZ construct. Before fixation, cells were synchronized in G2 phase with 9 µM RO3306 for 15 h and then released into mitosis. Subsequently, cells were immediately treated with 10 µM MG132 for an additional hour. CENP-C was used to visualize kinetochores and DAPI to stain DNA. Scale bar: 5 µm. (**B**) Quantification of Zwilch levels at kinetochores of the experiment shown in panel A. n refers to individually measured kinetochores. (**C**) Representative images showing cells treated as in panel A. (**D**) Quantification of Zwilch levels at kinetochores of the experiment shown in panel C. n refers to individually measured kinetochores. (**E**) Representative images showing the localization of the indicated $^{mCh}$RZZ constructs in prometaphase after depletion of Zwilch with 100 nM. 61 h after

Zwilch RNAi treatment cells were electroporated with mCherry or different $^{mCh}$RZZ constructs as indicated. Following an 8 h recovery, cells were synchronized in G2 phase with 9 µM RO3306 for 15 h and then released into mitosis. Subsequently, cells were immediately treated with 3.3 µM Nocodazole for an additional hour. CENP-C was used to visualize kinetochores and DAPI to stain DNA. Scale bar: 5 µm. (**F**) Quantification of Zwilch levels at kinetochores of the experiment shown in panel E. n refers to individually measured kinetochores. (**G**) Representative images showing the localization of the indicated $^{mCh}$RZZ constructs in prometaphase after depletion of CENP-E with 60 nM siRNA and Zwilch with 100 nM as shown in (Figure 5C). 13 h after CENP-E RNAi treatment cells were electroporated with different $^{mCh}$RZZ constructs as indicated. Following an 8 h recovery, cells were synchronized in G2 phase with 9 µM RO3306 for 15 h and then released into mitosis. Subsequently, cells were immediately treated with 3.3 µM Nocodazole, 10 µM MG132 and, where indicated, with 500 nM Reversine, for an additional hour. CENP-C was used to visualize kinetochores and DAPI to stain DNA. Scale bar: 5 µm. (**H-I**) Quantification of Zwilch levels at kinetochores of the experiment shown in panel G. n refers to individually measured kinetochores.

***Figure 5 – Supplement 2***

(**A**) Representative images showing kinetochore levels of Zwilch after depletion of Spindly and CENP-E. Spindly RNAi treatment was performed with 50 nM siRNA. 24 h after Spindly RNAi treatment, cells were transfected with 60 nM CENP-E siRNA. 8 h after transfection, cells were synchronized in G2 phase with 9 µM RO3306 for 15 h and then released into mitosis. Subsequently, cells were immediately treated with 3.3 µM Nocodazole, 10 µM MG132 and, where indicated, with 500 nM Reversine, for an additional hour. CENP-C was used to visualize kinetochores and DAPI to stain DNA. Scale bar: 5 µm. (**B**) Quantification of Zwilch levels at kinetochores of the experiment shown in panel A. n refers to individually measured kinetochores.

***Figure 6 – Supplement 1***

(**A**) Representative images of DLD-1 cells after depletion of KNL1 and Ndc80C. Ndc80C RNAi treatment was performed with two transfections of 20 nM siRNA directed against Hec1, SPC24, and SPC25 subunit. KNL1 RNAi was performed with 60 nM siRNA. 8 h after the second siNdc80C RNAi and KNL1 RNAi treatment, cells were synchronized in G2 phase with 9 µM RO3306 for 15 h and then released into mitosis. Subsequently, cells were immediately treated with

3.3 µM Nocodazole and 10 µM MG132 for an additional hour. CENP-C was used to visualize kinetochores and DAPI to stain DNA. Scale bar: 5 µm. (**B-E**) Quantification of Zwilch, CENP-E, KNL1, and Hec1 levels at kinetochores of the experiment shown in panel A. n refers to individually measured kinetochores. (**F**) Quantification of Zwilch levels at kinetochores of the experiment described in Figure 6A. (**G**) Representative images showing cells electroporated with $^{mCh}R^{EE}ZZ$. Ndc80C RNAi treatment was performed with two transfections of 20 nM siRNA directed against HEC1, SPC24, and SPC25 subunit. KNL1 and CENP-E RNAi were performed with 60 nM siRNA. 13 h after the second siNdc80C RNAi treatment and CENP-E or KNL1 RNAi treatment cells were electroporated with $^{mCh}R^{EE}ZZ$. Following an 8 h recovery, cells were synchronized in G2 phase with 9 µM RO3306 for 15 h and then released into mitosis. Subsequently, cells were immediately treated with 3.3 µM Nocodazole and 10 µM MG132 for an additional hour. CENP-C was used to visualize kinetochores and DAPI to stain DNA. Scale bar: 5 µm. (**H-I**) Quantification of the experiment shown in panel G. n refers to individually measured kinetochores. (**J-L**) Quantification of CENP-E, BUB1, and KNL1 levels at kinetochores of the experiment described in Figure 6D. n refers to individually measured kinetochores.

***Figure 6 – Supplement 2***

(**A**) Representative images of HeLa cells after depletion of Ndc80C and BUB1. Ndc80C RNAi treatment was performed with two transfections of 20 nM siRNA directed against Hec1, SPC24, and SPC25 subunit. BUB1 RNAi was performed with 50 nM siRNA. 8 h after the second siNdc80C RNAi and BUB1 RNAi treatment, cells were synchronized in G2 phase with 9 µM RO3306 for 15 h and then released into mitosis. Subsequently, cells were immediately treated with 3.3 µM Nocodazole, 10 µM MG132 and, where indicated, with 500 nM Reversine, for an additional hour. CENP-C was used to visualize kinetochores and DAPI to stain DNA. Scale bar: 5 µm. (**B-C**) Quantification of Zwilch and CENP-E levels at kinetochores of the experiment shown in panel A. n refers to individually measured kinetochores.

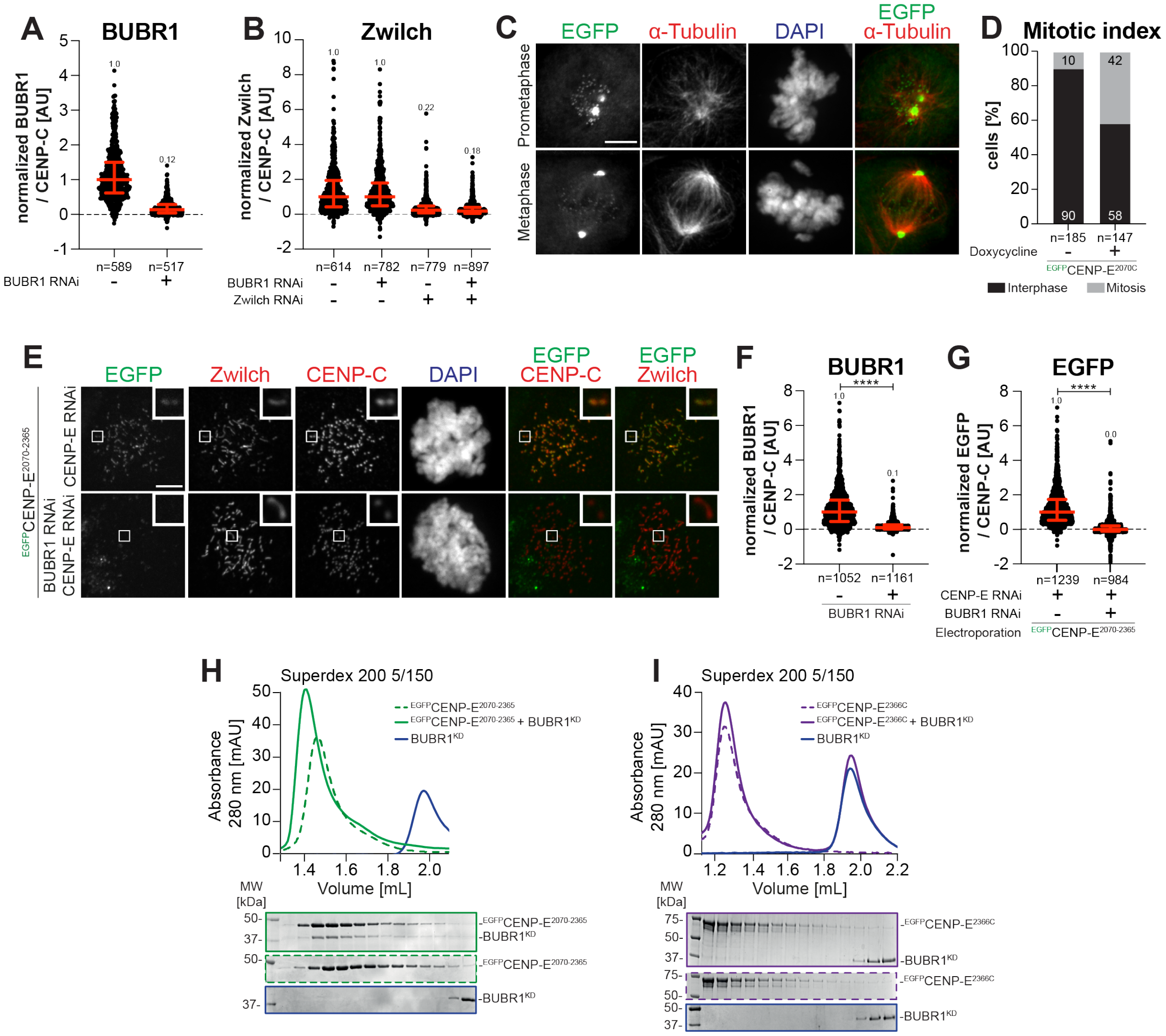


Figure 1
Supplement 1

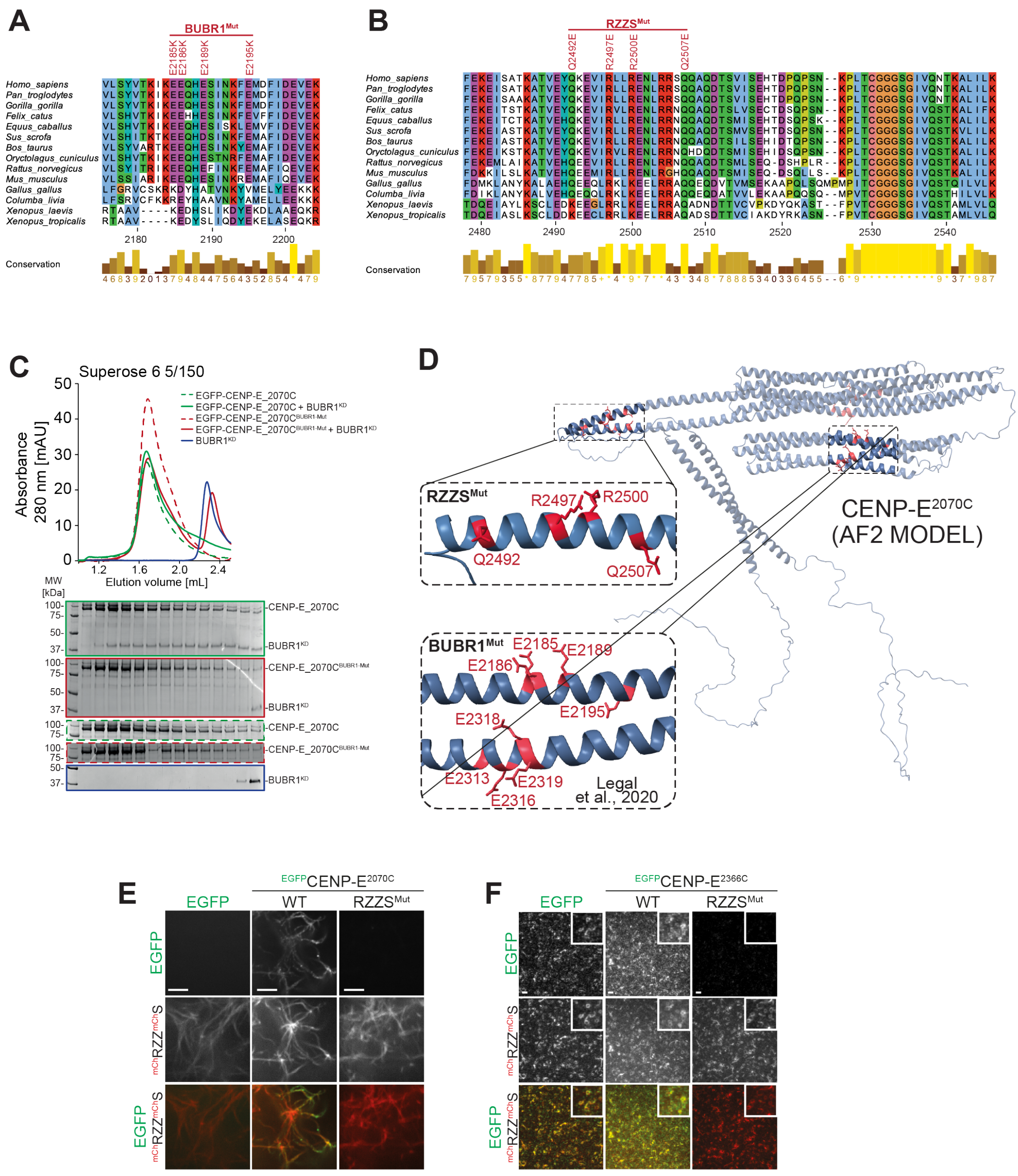


Figure 2
Supplement 1

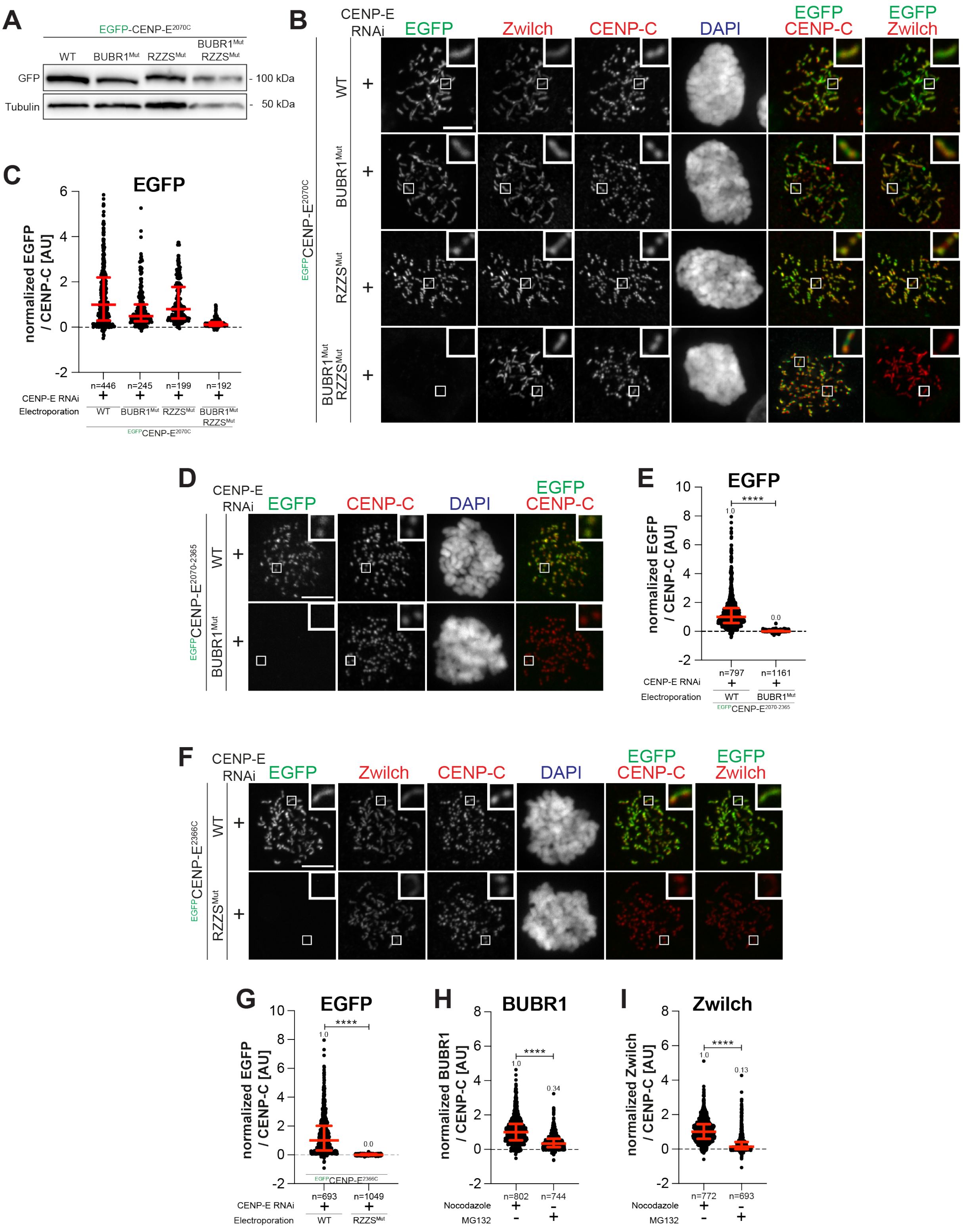


Figure 2
Supplement 2

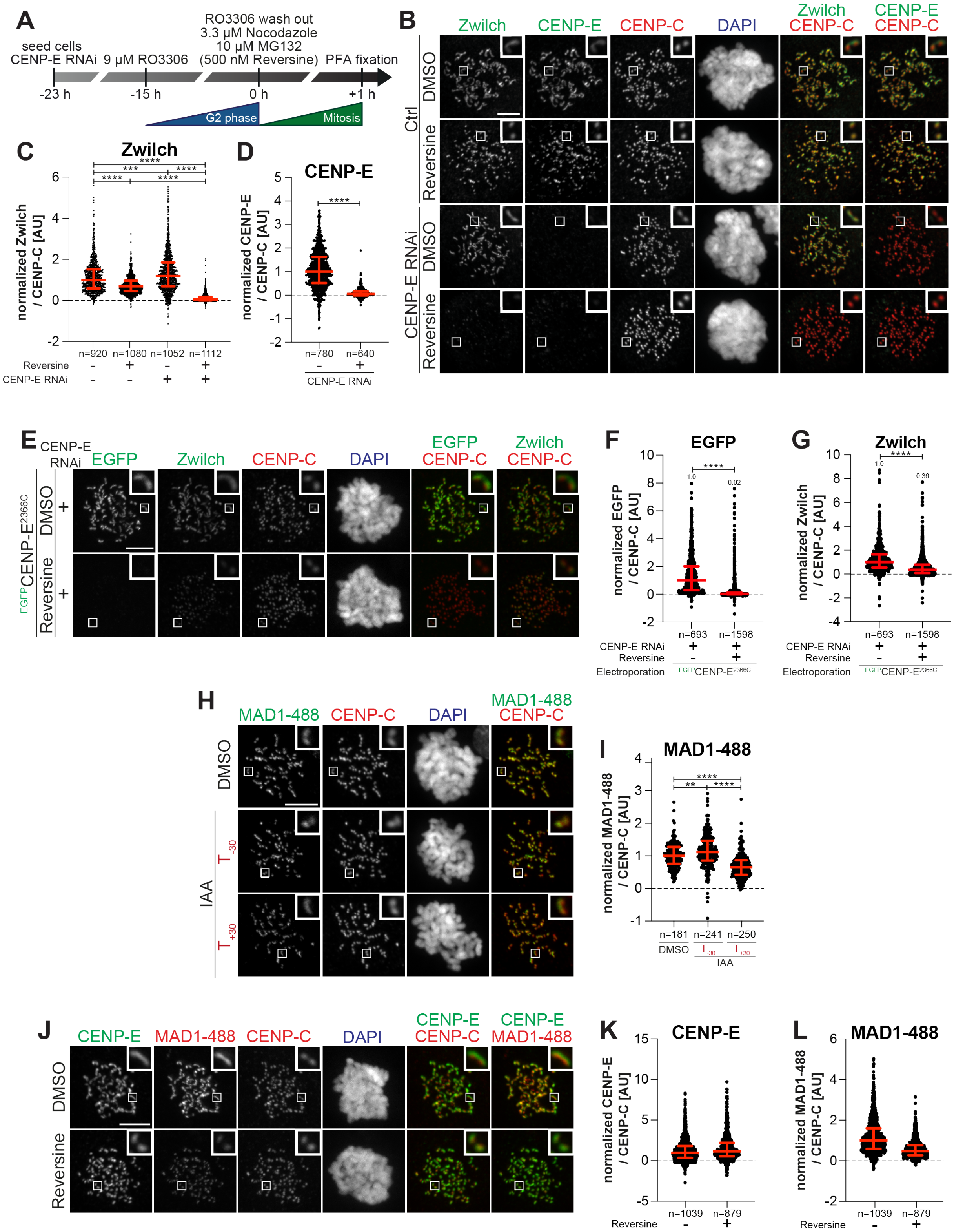


Figure 3
Supplement 1

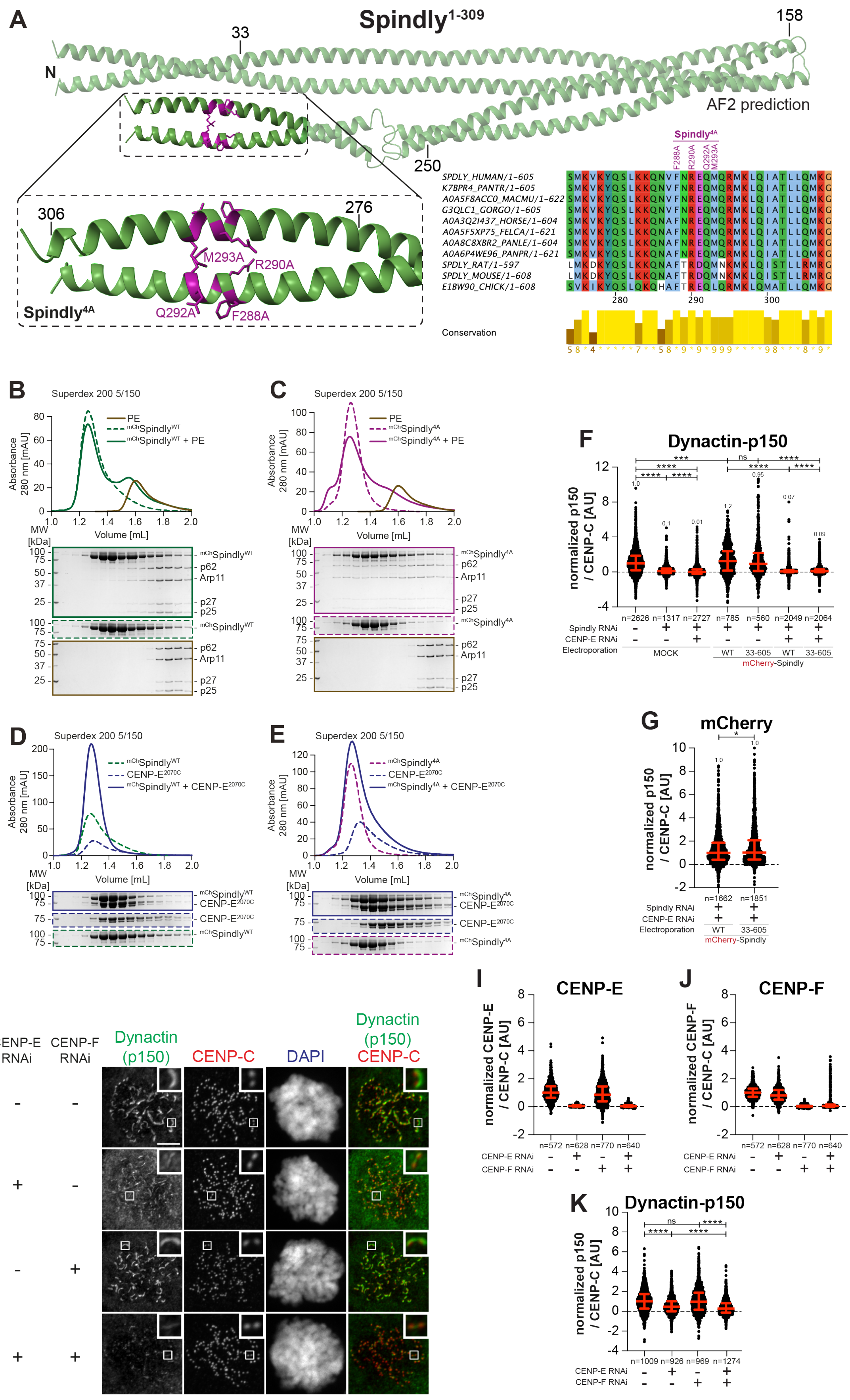


Figure 4
Supplement 1

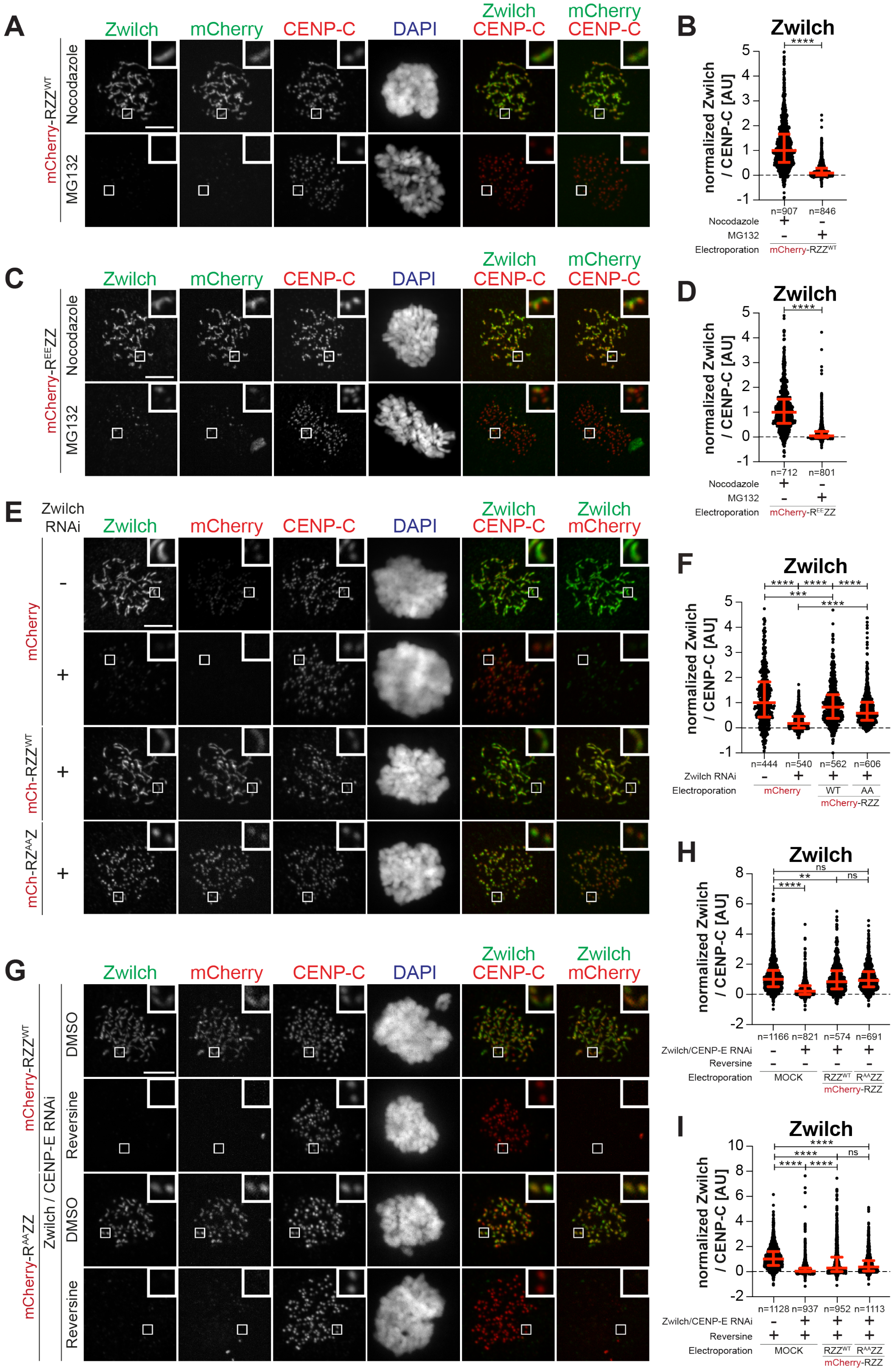


Figure 5
Supplement 1

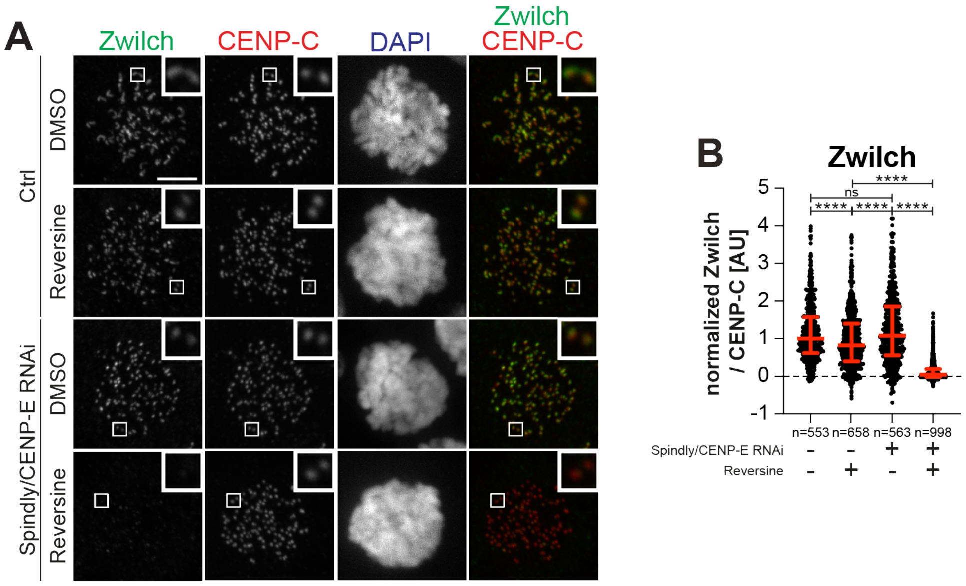


Figure 5 - Supplement 2

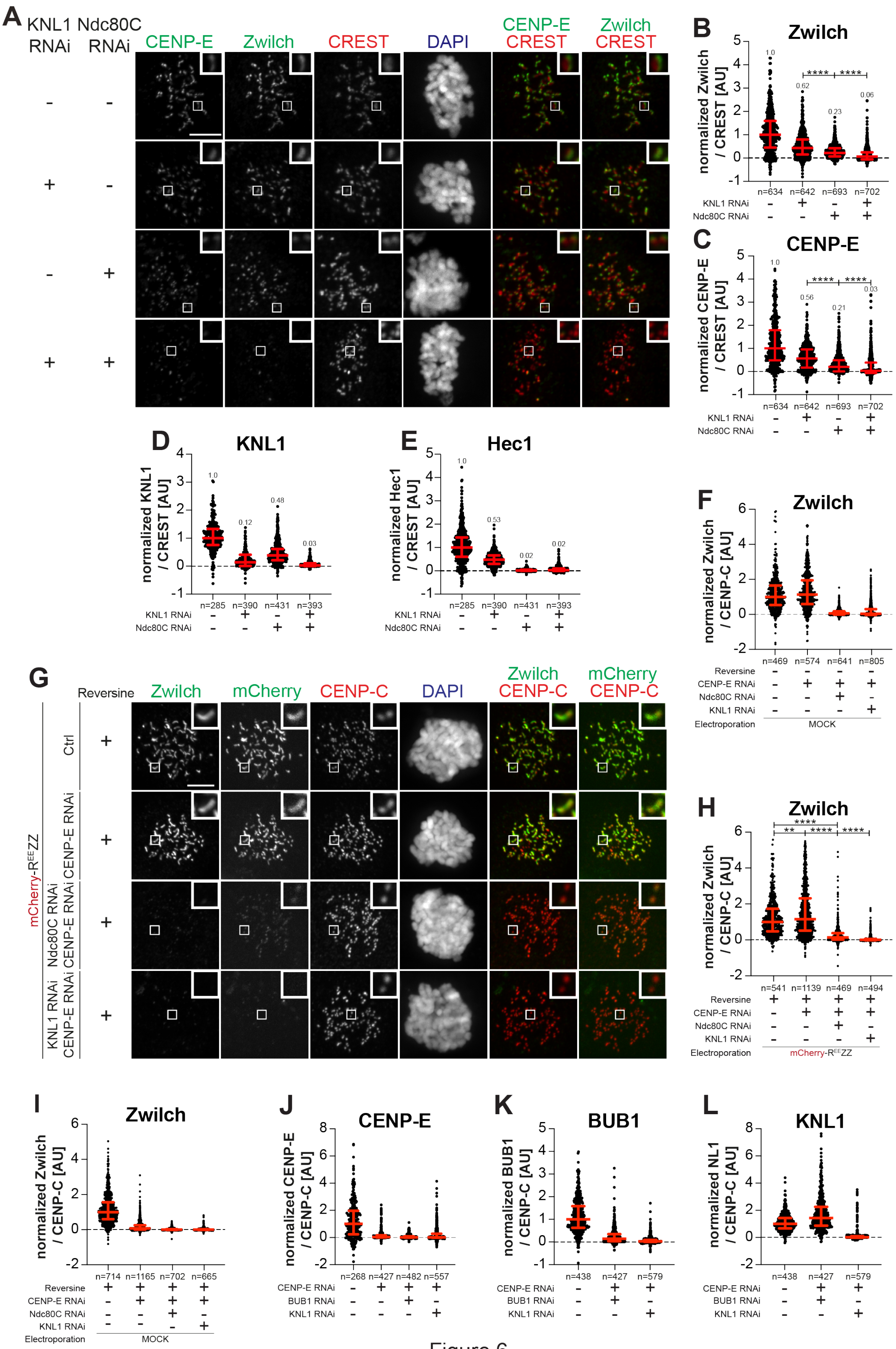


Figure 6
Supplement 1

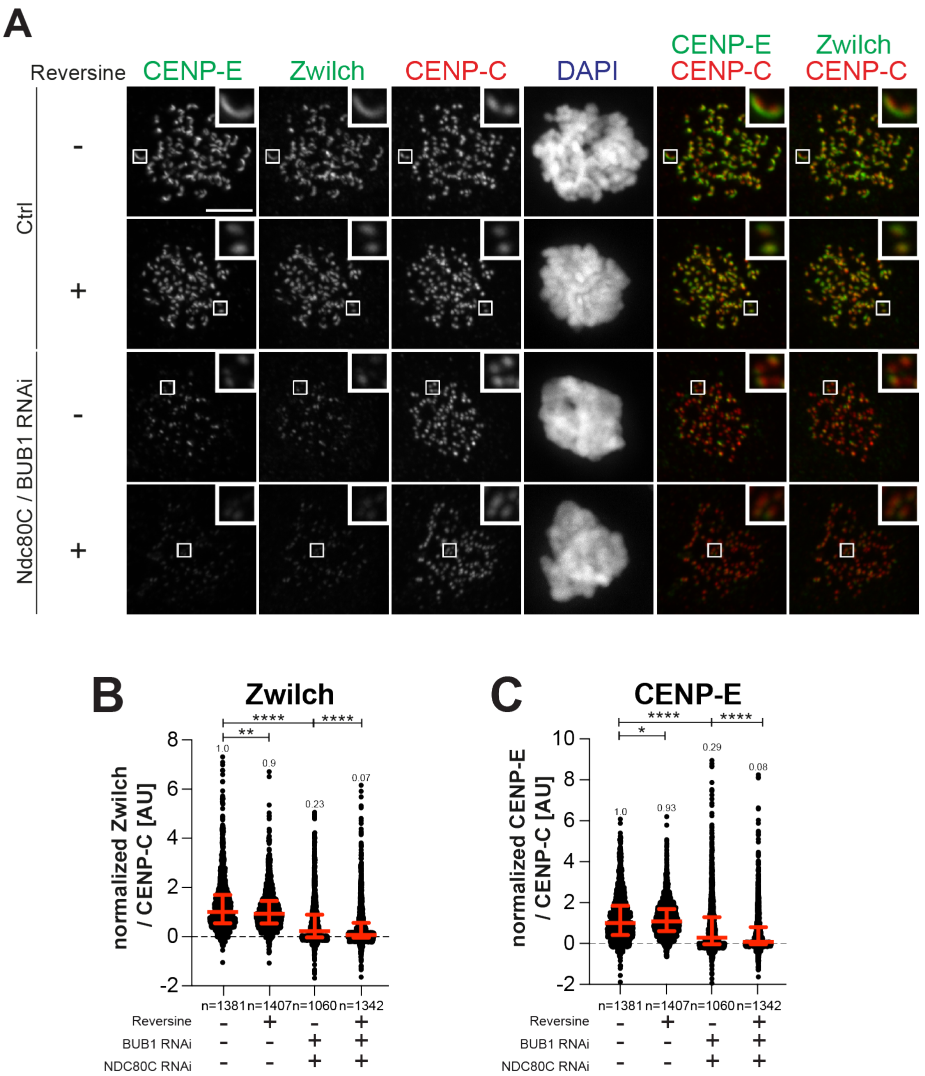


Figure 6
Supplement 2
